# A cell-penetrant peptide blocking *C9ORF72*-repeat RNA nuclear export suppresses neurodegeneration

**DOI:** 10.1101/2021.05.23.445325

**Authors:** Lydia M. Castelli, Alvaro Sanchez-Martinez, Ya-Hui Lin, Santosh Kumar Upadhyay, Adrian Higginbottom, Johnathan Cooper-Knock, Aytac Gül, Amy Walton, Claire Montmasson, Rebecca Cohen, Claudia S. Bauer, Kurt J. De Vos, Mimoun Azzouz, Pamela J. Shaw, Cyril Dominguez, Laura Ferraiuolo, Alexander J. Whitworth, Guillaume M. Hautbergue

**Affiliations:** Sheffield Institute for Translational Neuroscience (SITraN), University of Sheffield, 385a Glossop Road, Sheffield S10 2HQ, United Kingdom; MRC Mitochondrial Biology Unit, University of Cambridge, Cambridge Biomedical Campus, Hills Road, Cambridge CB2 0XY, United Kingdom; Leicester Institute of Structural & Chemical Biology and Department of Molecular & Cell Biology, University of Leicester, Leicester LE1 7RH, United Kingdom; INSERM U1270, Institut du Fer à Moulin, 75005 Paris, France; Manchester Centre for Genomic Medicine, Manchester University NHS Foundation Trust, M13 9WL, UK

**Author notes:** These authors contributed equally to this work (Lydia M. Castelli, Alvaro Sanchez-Martinez, Ya-Hui Lin).

## Abstract

Hexanucleotide repeat expansions in *C9ORF72* are the most common genetic cause of amyotrophic lateral sclerosis (ALS) and frontotemporal dementia (FTD), a spectrum of incurable debilitating neurodegenerative diseases. Here, we report a novel ALS/FTD drug concept with *in vivo* and *in vitro* therapeutic activity in preclinical models of C9ORF72-ALS/FTD. Our data demonstrate that supplementation or oral administration of a cell-penetrant peptide, which competes with the SRSF1:NXF1 interaction, confers neuroprotection by inhibiting the nuclear export of pathological *C9ORF72*-repeat transcripts in various models of disease including primary neurons, patient-derived motor neurons and *Drosophila*. Our drug-like rationale for disrupting the nuclear export of microsatellite repeat transcripts in neurological disorders provides a promising alternative to conventional small molecule inhibitors often limited by poor blood-brain barrier penetrance.

## Introduction

Polymorphic GGGGCC (G4C2) hexanucleotide repeat expansions in the *C9ORF72* gene cause the most common genetic forms of amyotrophic lateral sclerosis (ALS) and frontotemporal dementia (FTD) (*1, 2*). These fatal adult-onset neurodegenerative disorders, which can co-manifest in the same individuals or families, respectively lead to muscle weakness linked to progressive paralysis and altered cognitive and personality features associated with neurodegenerative changes in the frontal and temporal lobes of the brain (*3*). A growing body of evidence in cell and animal models pinpoints RNA gain-of-toxic functions through repeat-associated non-AUG (RAN) translation of dipeptide repeat proteins (DPRs) produced in all reading frames from sense and antisense transcripts as one of the main drivers of disease pathophysiology (*4*). In agreement with this, we recently showed that sequestration of SRSF1 (SR-rich splicing factor 1) on hexanucleotide-repeat RNA and interaction with NXF1 (nuclear export factor 1) triggers the nuclear export of sense and antisense *C9ORF72* repeat transcripts and subsequent RAN translation of cytotoxic DPRs (*5*). Concordantly, depletion of SRSF1 confers neuroprotection in C9ORF72-ALS patient-derived motor neurons and *Drosophila* (*5*).

No effective neuroprotective treatment is currently available for ALS or FTD patients. The standard of care involves treatment with riluzole, an anti-glutamate agent, for ALS (*6*) and serotonin reuptake inhibitors such as citalopram for FTD (*7*). However, none of these drugs treat neuronal loss and riluzole only extends the life of ALS patients by approximately 3 months. A Phase 1 clinical trial based on the intrathecal injection of antisense-oligonucleotides (BIIB078, *Biogen;* US NCT03626012, UK MND CSG) to downregulate the expression of pathological *C9ORF72*-repeat transcripts (*8*) is underway for C9ORF72-ALS patients. However, only pathological sense transcripts are targeted via this genetic therapy approach. Overall, there are few drugs available for neurodegenerative disorders largely due to poor blood-brain barrier penetrance, issues with targeting protein-protein interactions and incomplete knowledge of disease-causing mechanisms. Over 50 conventional drug-based clinical trials for ALS have failed in the past 20 years (*9*). Conversely, peptides of biological interest fused to a protein transduction domain (PTD) mediating intra-cellular delivery have emerged as potential therapeutic compounds in pre-clinical models of cerebral ischemia, Duchenne muscular dystrophy, cardiac diseases and oncology models (*10*). We and others have also shown that these cell permeable/penetrating peptides (CPPs) form useful molecular tools to specifically target biological pathways in mammalian cells (*11, 12*). Here, we evaluated the efficacy of SRSF1-based CPPs to potentially inhibit the SRSF1:NXF 1 interaction, which licenses the nuclear export of pathological *C9ORF72*-repeat transcripts, as a novel therapeutic concept in ALS/FTD.

## Results and Discussion

SRSF1 amino-acids (aa) 89-120 comprising the unstructured linker sequence between the two RNA recognition motifs (**Fig. S1A**) are required and sufficient for interaction with the aminoterminal RNA-binding region of NXF1 (aa 1-198, **Fig. S1B**) using pull-down assays with recombinant GST-SRSF1 fusions and ^35^S-methionine radiolabeled NXF1 protein domains (**Fig. S1C**). We previously showed that arginines 90, 93, 117 and 118 of SRSF1 mediate the interaction with NXF1 (*13*). To generate SRSF1-CPPs, either wild-type (WT) or 4 arginine-mutated (m4) linker regions of SRSF1 were fused to a V5 tag and the PTD of the HIV1 TAT protein (aa 47-57), together with a control (Ctrl)-CPP lacking the SRSF1 sequence (**Fig. 1A**). The experimental molar extinction coefficients of the cell permeable peptides were determined to further allow accurate quantification of CPPs (**Fig. S2**). High content immunofluorescence microscopy imaging showed that the CPPs are efficiently delivered and maintained within the nucleus and cytoplasm of >95% human V5-stained HEK293T cells when added to the cell culture medium for 72 hours at 1-10 μM (**Fig. 1B-C**). Importantly, SRSF1 WT-CPP specifically inhibits the co-immunoprecipitation of endogenous SRSF1 with FLAG-tagged NXF1, while co-interacting with NXF1 in HEK293T cells (**Fig. 1D**), indicating that WT-CPP disrupts the SRSF1:NXF1 interaction in human cells. To precisely characterize the molecular mechanism of action, we performed pull-down assays using bacterially-expressed proteins purified in high salt and in the presence of RNase to eliminate the detection of false positive interactions bridged by RNA molecules. These assays show that WT-CPP interacts directly with NXF1 in a dose-dependent manner (**Fig. 1E**) directly competing with the SRSF1:NXF1 interaction (**Fig. 1F**). As a specificity control, mutation of the 4 arginines involved in the interaction with NXF1 substantially reduced the binding of m4-CPP to NXF1 (**Fig. 1E**) and higher concentrations of m4-CPP were thus required to disrupt the SRSF1:NXF1 interaction (**Fig. 1F**). Isothermal titration calorimetry assays (ITC) further quantified the affinity of WT-CPP and of the minimal NXF1-interacting sequence of SRSF1 comprising aa89-120 (**Fig. S3A**) with the SRSF1/RNA-binding site of NXF1 (aa1-198, Fig. S1B-C). The heat released during binding to NXF1 is shown for WT-CPP and SRSF1 aa89-20 wild-type or SRSF1-m4 mutant peptide sequences in **Fig. S3B-D**. As expected from the above pull-down assays described above, transformation of the thermodynamic parameters acquired from the heat-released titration experiments into protein-ligand binding titration curves further confirmed that the mutated SRSF1-m4 sequence affinity for NXF1 is reduced approximately 10-fold compared to SRSF1 WT-CPP (**Table S1**). Interestingly, the affinity of SRSF1 WT-CPP compared to the wild-type SRSF1 peptide only is increased 5-fold, likely due to the presence of arginine residues in the TAT protein transduction domain, thus conferring an increased potency of the WT CPP drug-like compound for disrupting the SRSF1:NXF1 interaction. In addition, we also validated that WT-CPP competes with the interaction between full length mammalian SRSF1 synthesized in rabbit reticulocytes and recombinant NXF1 (**Fig. S4**). Taken together, these data demonstrate that SRSF1-CPPs are delivered intracellularly and inhibit the SRSF1:NXF1 interaction *in vivo* in human cells through direct competition.

**Fig. 1.**
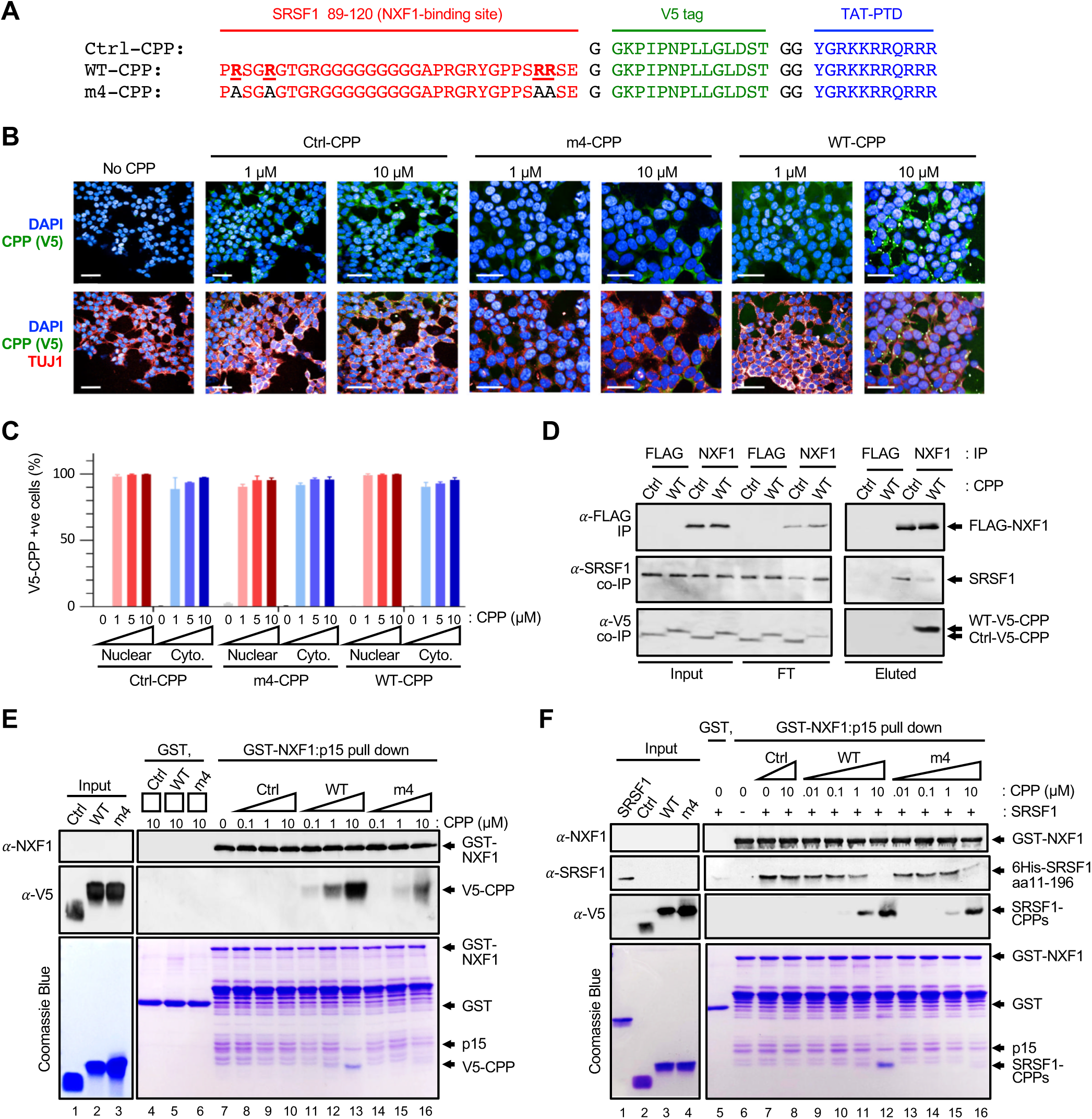
SRSF1-CPPs are intracellularly delivered and inhibit the interaction of endogenous SRSF1 with NXF1 through direct competition. (**A**) Cell permeable peptide sequences composed of SRSF1 aa89-120 bearing or not Arg-Ala substitutions of 4 arginines involved in the interaction with NXF1, a V5 tag and the TAT protein transduction domain (TAT-PTD). (**B**) High content immunofluorescence imaging microscopy of HEK293T cells cultured in media supplemented with Ctrl, m4 and WT CPPs for 72 hours. CPPs were detected in the green channel using anti-V5 staining. DAPI was used to delineate the nucleus while TUJ1 staining (in the red channel) was used to delineate the cytoplasm. Scale bar: 50 μm. (**C**) Quantification of V5-CPP positive HEK293T cells visualised in (B) was statistically assessed in a blinded automated manner from approximately 3000 cells cultured with each CPP. (**D**) Co-immunoprecipitation assays from HEK293T cells transfected for 72h with either FLAG control or FLAG-NXF1 expressing plasmids and either Ctrl-CPP or WT-CPP. 12% and 20% SDS-PAGE gels were run to respectively analyse the levels of NXF1 (71 kDa) or SRSF1 (28 kDa) and CPPs (Ctrl-CPP: 3 kDa; SRSF1-CPP: 6.1 kDa). (**E**) Pull down assay using recombinant GST and GST-NXF1:p15 fusions with increasing amounts of Ctrl-CPP, WT-CPP and m4-CPP. Note that GST-NXF1 was co-expressed in bacteria with p15/NXT1, a co-factor which improves the expression of NXF1 through heterodimerization with the NTF2-like domain (aa371-551) of NXF1 (*25, 26*). (**F**) Competitive binding assays using purified GST, GST-NXF1 and SRSF1 11-196 recombinant proteins expressed in *E. coli* together with increasing concentrations or Ctrl-CPP, WT-CPP or m4-CPP. Full length SRSF1 does not express in *E. coli* and could not be used here however Fig. S1C showed that aa89-120 of SRSF 1 constitute the NXF1 – binding site.

We next tested whether SRSF1 WT-CPP would inhibit the SRSF1:NXF1-dependent nuclear export and RAN translation of *C9ORF72*-repeat transcripts in human cell models of C9ORF72-ALS/FTD. To facilitate the quantification of RAN-translated DPRs, we generated uninterrupted repeat constructs lacking canonical AUG start codons and comprising 45 G4C2 or 43 C4G2 hexanucleotide repeats with downstream V5 tags in the 3 coding frames to allow detection of all DPRs species in a single western blot assay. Plasmid maps of the engineered sense and antisense reporter plasmids are respectively presented in **Figs. S5** and **S6**. Addition of increasing concentrations of WT-CPP to the medium of HEK293T cells transfected with G4C2_45_-3xV5 or C4G2_43_-3xV5 repeat transcript constructs led to dose-dependent inhibition of nuclear export with nuclear increase associated with concomitant cytoplasmic decrease of *C9ORF72*-repeat transcripts while Ctrl-CPP had no effect (**Fig. 2A-B**). Western blotting analysis of total, nuclear and cytoplasmic fractions validated the absence of nuclear contamination in the cytoplasmic samples probed with the chromatin-associated SSRP1 factor and the predominantly cytoplasmic HSPA14 heat shock protein (**Fig. S7A-B**). Interestingly, the SRSF1-CPP-induced nuclear export inhibition of *C9ORF72*-repeat transcripts correlates with a dose-dependent reduction of the RAN-translation of sense (poly-GA, GP, GR) and antisense (poly-GP, PR, PA) DPRs in the same conditions (**Fig. 2C-D**). SRSF1 m4-CPP also inhibited the production of sense and antisense DPRs, but to a lesser extent, in agreement with its reduced affinity for NXF1. As expected, quantification of DPR expression levels using western blots probed with a V5 antibody showed that SRSF1-CPPs lead to similar inhibitory dose-responses for sense and antisense DPRs (**Fig. 2 E-F**) with IC50 concentrations of approximately 0.5 μM for WT-CPP and 1.0 μM for m4-CPP (**Fig. 2G-H**). Expression of sense and antisense DPRs from the G4C2_45_-3xV5 and C4G2_43_-3xV5 repeat transcript constructs into HEK293T cells transfected in the above conditions is further correlated with reduced cell proliferation as measured in MTT assays while 0.5 μM WT-CPP and 1.0 μM m4-CPP respectively rescue the DPR-associated cytotoxicity in the C9ORF72-ALS/FTD cell models (**Fig. 2I-J**). Taken together, our data show that SRSF1-CPPs efficiently inhibit the nuclear export and subsequent RAN translation of both sense and antisense reporter *C9ORF72*-repeat transcripts in human HEK293T cell models of ALS/FTD.

**Fig. 2.**
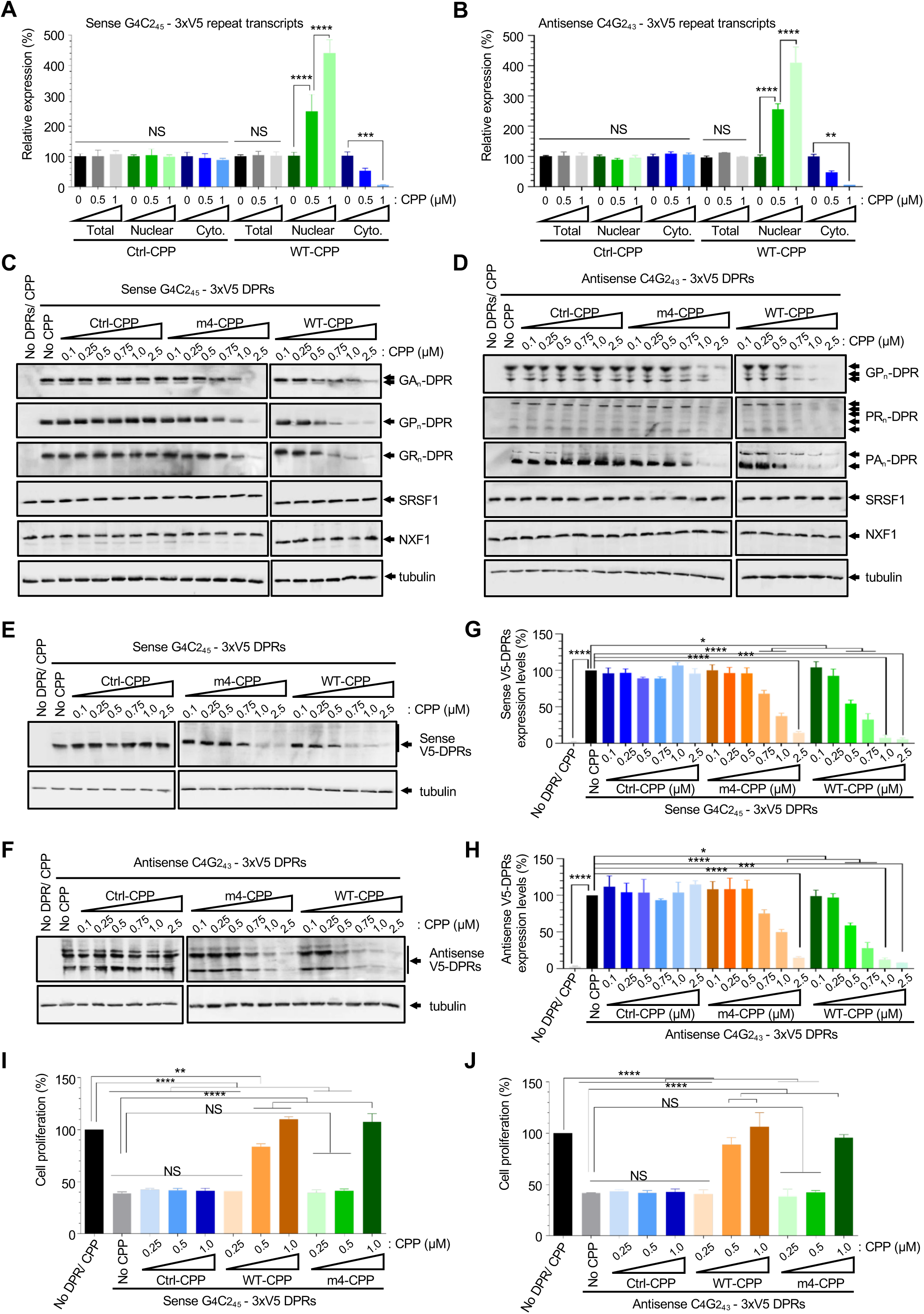
SRSF1 WT-CPPs inhibits the nuclear export of reporter *C9ORF72*-repeat transcripts rescuing the DPR-associated cytotoxicity in human cell models of C9ORF72-ALS. (**A**) Total, nuclear and cytoplasmic levels of sense G4C2_45_-3xV5 repeat transcripts from 72h-transfected cells were quantified by qRT-PCR in biological triplicate experiments following normalization to *U1* snRNA levels and to 100% in untreated cells (mean ± SEM; one-way ANOVA with Tukey’s correction for multiple comparisons, NS: non-significant, *: p<0.05, ***: p<0.001, ****: p<0.0001). (**B**) Total, nuclear and cytoplasmic levels of antisense C4G2_43_-3xV5 repeat transcripts from 72h-transfected cells were quantified by qRT-PCR in three biological triplicates following normalization to *U1* snRNA levels and to 100% in untreated cells (mean ± SEM; one-way ANOVA with Tukey’s correction for multiple comparisons, NS: non-significant, *: p<0.05, ***: p<0.001, ****: p<0.0001). (**C, D**) Western blot analysis from cells transfected for 72h with no DPR control or G4C2_45_-3xV5 (C) and C4G2_43_-3xV5 (D) plasmids for 6 hours prior to medium change with media supplemented with increasing concentrations of Ctrl, m4 or WT CPPs. Blots are probed for poly GR, GP and GA DPRs along with SRSF1, NXF1 and tubulin. (**E, F**) Western blots from cells transfected with no DPR control or G4C2_45_-3xV5 and C4G2_43_-3xV5 plasmids and treated with Ctrl, m4 or WT CPPs as in (C, D). Blots are probed for V5 and !-tubulin. (**G**) Intensities of V5-tagged sense DPR bands in panel E were quantified relative to their !-tubulin loading control levels in biological triplicates (Mean ± SEM; one-way ANOVA; *: p<0.05, ***: p<0.001, ****:p<0.0001). (**H**) Intensities of V5-tagged antisense DPR bands in panel F were quantified relative to their !-tubulin loading control levels in biological triplicates (Mean ± SEM; one-way ANOVA; *: p<0.05, ****:p<0.0001). (**I-J**) MTT cell proliferation assays from cells transfected and treated with CPPs as in panels C-H were performed in biological triplicates (mean ± SEM; one-way ANOVA, **: p<0.01, ***: p<0.001, ****: p<0.0001).

To investigate the relevance of these findings in neurons, high content V5-immunofluorescence imaging microscopy showed that the CPPs are also efficiently delivered and maintained into >90% primary rat neurons when added to the medium for 72 hours at 5-10 μM, with >80% neurons showing detectable levels of SRSF1-CPPs at 1 μM (**Fig. 3A-B**). We next generated lentivirus (LV) expressing the G4C2_45_-3xV5 or C4G2_43_-3xV5 repeat transcripts and transduced them for 16 hours into the cortical neuron cultures prior to replacing the media with fresh medium supplemented with either Ctrl-CPP or SRSF1-CPPs. Western blot analysis of V5-tagged DPRs conducted 72 hours post CPP addition showed that WT-CPP, and m4-CPP to a lesser extent, efficiently inhibited the production of sense and antisense DPRs with a pronounced effect at higher concentrations (**Fig. 3C**). Consistent with this, higher concentrations of CPPs led to increased intracellular delivery of V5-tagged CPPs in the neuronal extracts loaded for western blot analysis. Lentiviral-mediated depletion of SRSF1 was further used as a positive control (*5*) showing that both depletion of SRSF1 or addition of 5 μM WT-CPP similarly inhibited the expression of sense and antisense DPRs (**Fig. 3C**). V5-immunofluorescence microscopy staining of DPRs in rat cortical neurons cultured in the same conditions but treated with 5 μM of Ctrl-CPP or WT-CPP synthesized without the V5 tags corroborated the previous data, showing that WT-CPP leads to a reduction in both the intensity and the number of neurons with V5-DPR signal (**Fig. S8A**). Blinded quantification further showed significant reduction in both sense and antisense V5-DPRs positive neurons treated with either LV-SRSF1-RNAi or WT-CPP (**Fig. S8B**), despite underestimation of the SRSF1 depletion or WT-CPP effects since neurons with successful SRSF1-targeted reduction levels of DPRs remain counted as DPRs-positive cells.

**Fig. 3.**
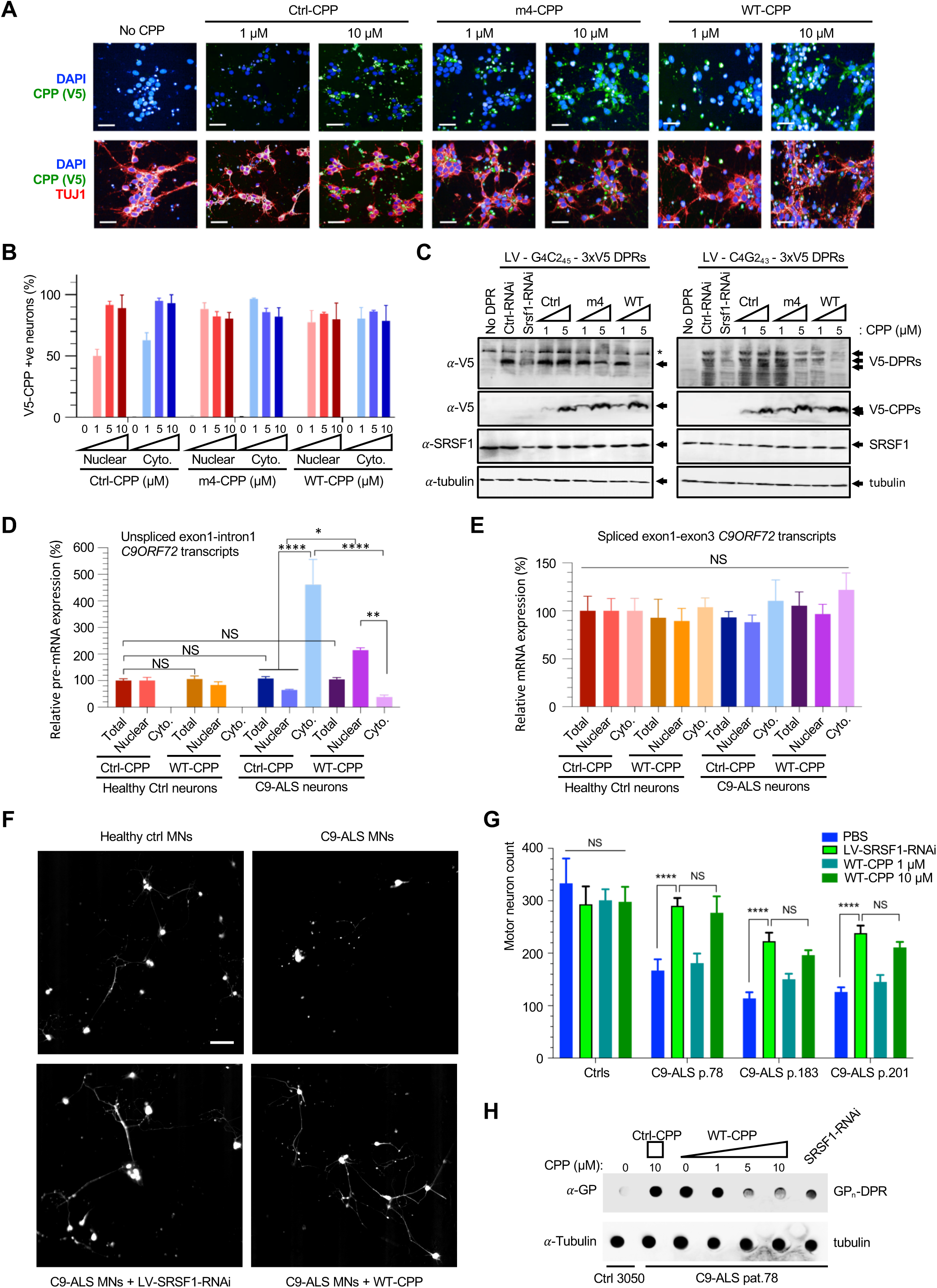
SRSF1 WT-CPP confers neuroprotection and inhibits RAN-translation of DPRs in C9ORF72-ALS primary neuron models and patient-derived motor neurons. (**A**) High content immunofluorescence imaging microscopy of primary rat cortical neurons cultured in media supplemented with Ctrl, m4 and WT CPPs for 72 hours. CPPs were detected in the green channel using anti-V5 staining. DAPI was used to delineate the nucleus while TUJ1 staining (in the red channel) was used to delineate the cytoplasm. Scale bar: 50 μm. (**B**) Quantification of V5-CPP positive neurons visualised in panel A was statistically assessed in a blinded automated manner from approximately 3000 cells cultured with each CPP. (**C**) Western blots from cultured rat cortical neurons transduced for 16 hours with G4C2_45_-3xV5, C4G2_43_-3xV5, Ctrl-RNAi or SRSF1-RNAi expressing lentivirus were treated for 72h with Ctrl, m4 and WT CPPs. Blots run on 12%SDS-PAGE are probed for V5 antibody to detect the V5-tagged DPRs (17-35 kDa), SRSF1 (28 kDa) and !-tubulin (52 kDa). 20% SDS-PAGE gels were run to detect V5-tagged CPPs (Ctrl-CPP: 3 kDa; SRSF1-CPPs: 6.1 kDa). * denotes a non-specific cross-reaction of the antibody. (**D-E**) qRT-PCR quantification of total, nuclear and cytoplasmic levels of pathological *C9ORF72*-repeat transcripts (D) and of intron-1-spliced *C9ORF72* transcripts (E) in 3 lines of patient-derived iNeurons (mean ± SEM; one-way ANOVA with Tukey’s correction for multiple comparisons, NS: non-significant, *: p<0.05, **: p<0.01, ****: p<0.0001). (**F**) High content imaging analysis of healthy control and C9ORF72-ALS iMotor Neuron survival in co-cultures with control or C9ORF72-ALS iAstrocytes. For lentiviral-mediated depletion of SRSF1, iAstrocyes and iMNs were separately transduced with LV-SRSF1-RNAi 48 hours prior to co-cultures for a period of 72 hours. CPPs were added to the medium of the co-cultures for 72 hours. See Supplementary Materials for a detailed timeline and protocol. Scale bar: 50 μm. (**G**) Automated quantification of motor neuron survival after 72 hours in co-cultures with iAstrocytes in independent triplicate lines (iMN count ± SEM; two-way ANOVA with Tukey’s correction for multiple comparisons, NS: non-significant, ****: p<0.0001). Data from 3 different healthy control cell lines were pooled together. (**H**) Dot blot assay probed for the poly-GP DPRs in Ctrl and C9ORF72-ALS patient-derived motor neurons treated with Ctrl-CPP, WT-CPP or SRSF1-RNAi expressing lentivirus. DPRs could only be detected in motor neurons derived from patient line 78 due to variability and low levels of DPRs across patient-derived lines.

The potential neuroprotective effects of SRSF1 WT-CPP was further tested in C9ORF72-ALS patient-derived cells. Fibroblasts from control individuals and C9ORF72-ALS patients (**Table S2**) were reprogrammed into induced neuronal progenitor cells that can be differentiated into induced-astrocytes or neurons and motor neurons (iAstrocytes, iNeurons, iMNs) (*5, 14, 15*). Advantageously, the differentiation of iNeurons yields a high number of cells allowing cellular fractionation experiments. Similar to the lentiviral-mediated depletion of SRSF 1 which inhibits the nuclear export of *C9ORF72*-repeat transcripts (*5*), 72-hour addition of 5 μM WT-CPP led to efficient block in iNeuron cultures showing increased nuclear and concomitant cytoplasmic reduction in the levels of *C9ORF72* transcripts retaining pathological repeat expansions in intron-1 (quantified by qRT-PCR using forward and reverse primers annealing in exon-1 and intron-1, **Fig. 3D**); while it had no effect on the expression of intron-1-spliced wild-type *C9ORF72* transcripts (measured with primers annealing in exon-1 and exon-3, **Fig. 3E**). The quality of the nuclear and cytoplasmic fractionation was validated in **Fig. S9**. Moreover, as for the depletion of SRSF1 in either iMNs or iAstrocytes (*5*), 72-hour addition of 5 μM WT-CPP promotes the survival of C9ORF72-ALS iMNs in co-cultures with C9ORF72-ALS iAstrocytes whilst it has no detrimental cytotoxic effects on heathy motor neurons (**Fig. 3F-G**). Consistent with the SRSF1-targeted nuclear export inhibition of repeat transcripts, both lentiviral-mediated depletion of SRSF 1 and addition of SRSF1 WT-CPP also lead to inhibition of the RAN translation of poly-GP DPRs encoded by both sense and antisense *C9ORF72*-repeat transcripts in mono-cultures of C9ORF72-ALS iMNs (**Fig. 3H**). Taken together, our data show that SRSF1 WT-CPP inhibits the nuclear export and subsequent RAN translation of DPRs, conferring neuroprotection in both C9ORF72-ALS patient-derived motor neurons and a primary neuronal model of disease.

We next sought to test the effects of CPPs *in vivo* in a *Drosophila* model of C9ORF72-ALS which expresses 36 uninterrupted G4C2 repeat transcripts and sense DPRs (*16*) by feeding flies with food supplemented with 10 μM CPPs. Whole-fly, nuclear and cytoplasmic fractions prepared from untreated or CPP-treated animals were initially verified showing absence of Histone H3 nuclear contamination in the cytoplasmic fractions and absence of cytoplasmic tubulin in the nuclear samples (**Fig. 4A**). Oral administration of SRSF1 WT-CPP led to specific reduction in the levels of cytoplasmic G4C236 repeat transcripts normalized to their total levels while m4-CPP showed a less pronounced non-statistically significant effect (**Fig. 4B**) indicating that similarly to the cell models, WT-CPP inhibits the nuclear export of *C9ORF72*-repeat transcripts in a *Drosophila* model of disease. Strikingly, this correlated with a reduction of the neurodegeneration-associated motor deficits observed in C9ORF72-ALS larvae (**Fig. 4C**) and adult flies (**Fig. 4D**) as well as an inhibition of the RAN translation of poly-GA, GP, GR DPRs (**Fig. 4E**). Ctrl-CPP and m4-CPP however show no significant effects on the rescue of locomotion or the expression of DPRs. Interestingly, both Ctrl-CPP and SRSF1-CPPs had no detrimental effects on the behavior of control healthy flies bearing 3 G4C2 repeats (**Figs 4C-D**) indicating that the CPPs are also safe to use and well-tolerated *in vivo*. Taken together, our data demonstrate that oral administration of SRSF1 WT-CPP mitigates the DPR-associated locomotor deficits in a C9ORF72-ALS *Drosophila* model of disease.

**Fig. 4.**
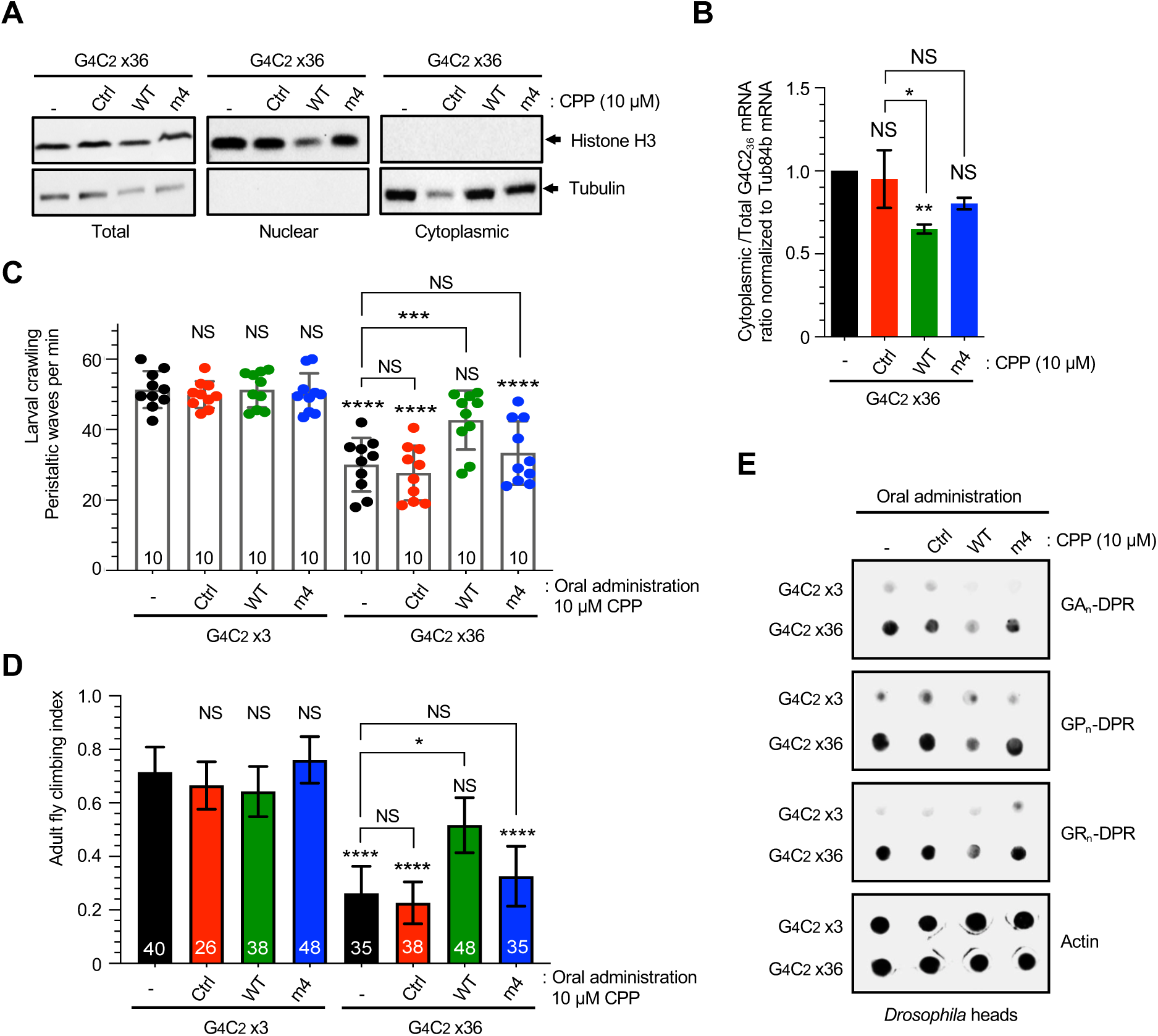
SRSF1 WT-CPP mitigates neurodegeneration-associated locomotor deficits and inhibits RAN-translation of DPRs in C9ORF72-ALS *Drosophila*. (**A**) Western blots of total, nuclear and cytoplasmic fractions isolated from G4C236 third instar larvae treated or not with Ctrl, WT and m4 CPPs were probed for nuclear Histone H3 and cytoplasmic tubulin proteins. Genotype: *da*-GAL4>*G4C2_36_*. (**B**) Total and cytoplasmic levels of sense G4C2_36_ repeat transcripts from G4C2_36_ third instar larvae treated or not with Ctrl, WT and m4 CPPs were quantified by qRT-PCR in biological triplicate experiments following normalization to Tub84b mRNA levels. Plot represents normalized Cytoplasmic/Total levels (mean ± SEM; one-way ANOVA with Sidak’s correction for multiple comparisons, NS: non-significant, *: p<0.05, **: p<0.01). Genotypes: *da*-GAL4>*G4C2_36_*. (**C**) Crawling assays in G4C23 healthy control or G4C236 third instar larvae fed or not with 10 μM Ctrl, WT or m4 CPPs (mean ± SEM; one-way ANOVA with Bonferroni’s correction for multiple comparisons, NS: non-significant, ***: p<0.001, ****: p<0.0001). Numbers of animals used in the assays are indicated at the bottom of each bar. Genotype: *nSyb*-GAL4>G4C2_3_, *nSyb*-GAL4>*G4C2_36_*. (**D**) Climbing assays in G4C2_3_ healthy control or G4C2_36_ 2 days old adult *Drosophila* fed or not with 10 μM Ctrl, WT or m4 CPPs (mean ± 95% CI normalized to control; Kruskal-Wallis test with Dunn’s correction for multiple comparisons, NS: nonsignificant, *: p<0.05, ****: p<0.0001). Numbers of animals used in the assays are indicated at the bottom of each bar. Genotypes: *D42*-GAL4>*G4C2_3_, D42*-GAL4>*G4C2_36_*. (**E**) Dot blot assays from G4C2_3_ healthy control or G4C2_36_ 2-day-old adult *Drosophila* heads fed or not with Ctrl, WT or m4 CPPs were probed for poly GA, GP and GR DPRs as well as for the loading control actin. Genotypes: *D42*-GAL4>*G4C2_3_, D42*-GAL4>*G4C2_36_*.

In this study, we report the use of a novel drug-like rationale to target the nuclear export of pathological microsatellite repeat transcripts through treatment with a cell-penetrant peptide which inhibits the SRSF1:NXF1-dependent nuclear export of *C9ORF72*-repeat transcripts and confers neuroprotection in preclinical patient-derived and *Drosophila* models of ALS/FTD. Moreover, no adverse effects of the CPPs were observed in control healthy patient-derived motor neurons/astrocytes or *Drosophila*. This study also shows that the canonical NXF1-dependent nuclear export of cellular mRNAs reviewed in (*17*) can be therapeutically targeted in *in vitro* and *in vivo* neurodegenerative models based on the inhibition of protein:protein interactions between mRNA nuclear export proteins. The neuroprotective effects conferred by oral administration of SRSF1 WT-CPP indicate efficient brain delivery. Previous studies have reported that TAT-based CPPs are able to cross the blood-brain barriers and be delivered to the central nervous system in mice (*18, 19*). Interestingly, oral administration of a poly-Q binding peptide 1 (QBP1)-based CPP, which interacts with poly-glutamine stretches and inhibits the aggregation of poly-Q proteins, suppressed neurodegeneration-associated premature death and inclusion body formation in a poly-Q *Drosophila* model (*20*) while improving the poly-Q phenotype of a mouse model through intra-cerebroventricular injection (*21*). Similarly, a cell permeable peptide bearing an RNA-binding motif of nucleolin, which binds poly-Q-encoded CAG-repeat RNA and sense *C9ORF72*-repeat reporter transcripts, rescued the rough eye neurodegenerative phenotypes in a *Drosophila* model of disease (*22, 23*). Our study further demonstrates successful intracellular delivery and efficacy of the SRSF1 cell permeable peptide in primary neurons and C9ORF72-ALS patient-derived neurons which harbor large pathophysiological expansions. Taken together, this highlights promising prospects for the development of CPP as drug-like compounds providing alternatives to conventional small molecule inhibitors which frequently have poor blood-brain barrier penetrance. Moreover, the development of CPPs for disrupting the nuclear export of other microsatellite repeat transcripts, which are involved in over 40 neurological disorders (*24*), could provide future promising agents for dissecting the nature of the pathogenic mechanisms linked to the formation of intranuclear RNA foci and RAN translation of polymeric repeat proteins as well as potential novel therapeutic strategies.

## Acknowledgments

We acknowledge Dr Adrian Isaacs (University College London, UK) for kindly providing the G4C2×3 and G4C2×36 *Drosophila* lines. This work was initially primed by the MND Association grant Hautbergue/Apr16/846-791 (GMH, LF, AJW, PJS) and is currently supported by the Biotechnology and Biological Sciences Research Council (BBSRC) grants BB/S005277/1 (GMH) and BB/S005579/1 (CD). GMH also acknowledges intermediate and current support from the Medical Research Council (MRC) New Investigator research grant MR/R024162/1. AJW was supported by MRC core funding (MC_UU_00015/6) and ERC Starting grant (DYNAMITO; 309742). LF was funded by the Thierry Latran Foundation (FTLAAP2016/ Astrocyte secretome) and the Academy of Medical Sciences Springboard Award. She is currently funded by the MRC grant MR/W00416X/1. KJDV acknowledges support from MRC grants MR/M013251/1 and MR/S025979/1. MA acknowledges support from from Alzheimer’s Research UK (ARUK-PG2018B-005), European Research Council (ERC Advanced Award 294745) and MRC DPFS (129016) grants. PJS is supported as an NIHR Senior Investigator (NF-SI-0617-10077), and by the MND Association (AMBRoSIA MNDAOct15/972-797) and the MRC Centres of Excellence for Neurodegeneration Pathfinder award (MR/SOO4920/1). We acknowledge ALS patients and their families for participating in our research and donating biological samples to the NIHR Sheffield Biomedical Research Centre: Translational Neuroscience for chronic neurological disorders (IS-BRC-1215-20017).

## Author contributions

GMH designed the overall study. GMH, AJW and CD respectively designed the cellular, *Drosophila* and biophysical assays. LF provided training for the differentiation of patient-derived neurons and did the co-culture assays. LMC and ASM respectively performed the molecular/cellular and *Drosophila* studies. LMC and YHL performed the nuclear export assays in patient-derived neurons. YHL and AG performed the biochemical studies. SKU performed the ITC assays. JCK, CM, RC and CSB were involved in the primary neuron experiments. AH was involved in the high content imaging. AW provided the dot blot. PJS provided genetically characterized human biosamples. GMH wrote the manuscript. LMC, ASM, YHL, PJS, CD, LF and AJW edited the manuscript. All authors approved the manuscript.

## Data and materials availability

All data are available in the main text or the supplementary materials. Plasmids generated in this study can be provided on request under a University of Sheffield Material Transfer Agreement.

## Supplementary Materials for

### Materials and Methods

#### Tissue culture

Cells were maintained in a 37 °C incubator with 5% CO_2_. HEK293T cells were cultured in Dulbecco’s Modified Eagle Medium (Lonza) supplemented with 10% fetal bovine serum (FBS) (Biosera) and 5 U ml^−1^ Penstrep (Lonza).

Primary rat cortical neurons were isolated at embryonic stage E18 from Sprague Dawley rat embryos (*Charles River*) and cultured and cultured on glass coverslips coated with poly-L-lysine in 12-or 24-well plates in neurobasal medium supplemented with B27 supplement (*Invitrogen*), 100 U/ml penicillin, 100 μg/ml streptomycin, and 2 mM L-glutamine. Cortical neurons were transfected using Lipofectamine LTX with PLUS reagent according to the manufacturer’s instructions (Thermofisher). Briefly, neurons were placed in fresh culture medium with a DNA:PLUS:LTX ratio of 1:0.5:0.5 (0.5–1 l μg DNA/100,000 cells/cm^2^). The transfection mix was replaced with conditioned medium after 6 hours. For lentiviral transduction, neurons were exposed to 4 MOI in fresh culture medium overnight (16 hours) prior to replacement with fresh medium containing CPPs for 72h.

For patient-derived cell cultures, informed consent was obtained from all subjects before sample collection (Study number STH16573, Research Ethics Committee reference 12/YH/0330). Human patient derived iAstrocytes, iNeurons and iMotor Neurons were differentiated from induced Neural Progenitor Cells (iNPCs) as described previously (*5, 14, 15*). Protocols of differentiation are also described below in the nuclear/cytoplasmic fractionation and co-culture assays.

#### Plasmids and cell permeable peptides (CPPs)

pCDNA3.1 plasmids bearing uninterrupted C9ORF72 hexanucleotide sense (GGGGCCx45) and antisense (CCCCGGx43) repeats were generated using annealed and concatemerized G4C2×15 Mung bean-blunted oligonucleotides cloned into the Klenow filled-in EcoRI site of pcDNA3.1-myc-HisA (*Invitrogen*) as described in (*5*). A 3x V5 tag cassette with V5 and stop codons in the 3 frames was custom synthesised (ThermoFisher Scientific) and subcloned downstream of the repeat sequences using the NotI/Xba1 sites to allow detection of DPR proteins in all three reading frames (Figs. S5 and S6). The hexanucleotide sense and antisense repeats with 3x V5 tags were subsequently subcloned from the pcDNA3.1 plasmids into the lentiviral plasmid SIN-PGK-cPPT-GDNF-WHV (*27*) using the BamHI and XhoI restriction sites. Lentiviral control and SRSF1-miRNA constructs were generated in (*5*). SRSF1 (amino-acids 11-196) was cloned into pET24b harbouring a 6His C-terminal tag (*13*). p3XFLAG-NXF1/TAP was generated in (*12*) using p3XFLAG-myc-CMV™-26 (*Sigma*). Full length TAP/NXF1 was amplified as a BamHI/XhoI PCR fragment and cloned into pGEX6P1 BamHI/XhoI (this study) prior to cotransformation into bacteria with pET9a-p15 (*12*) for recombinant purification of the GST-NXF1:p15 complex as in (*12*). All cell permeable peptides used in this study were custom synthesised by *ThermoFisher Scientific* at >90% purity (10-14 mg scale) and resuspended in PBS at 1 mM prior to flash freezing of aliquots in liquid nitrogen and storage at −80 °C. Design and sequence information is provided in Fig. 1A.

#### Lentivirus production

Lentivirus production was performed as described previously (Hautbergue et al 2017). Briefly, twenty 10 cm dishes seeded with 3 × 10^6^ HEK293T cells/dish were each transfected with 13 μg pCMVΔR8.92, 3.75 μg pM2G, 3 μg pRSV and 13 μg SIN-CMV-Ctrl-miRNA, SIN-CMV-SRSF1-miRNA, SIN-CMV-G4C2x45-3xV5 or SIN-CMV-C4G2x43-3xV5 using calcium phosphate transfection. Media was replaced after 12 h. After a further 48 h, the supernatant was collected, filtered through a 0.45 μm filter and centrifuged at 19,000 r.p.m. for 90 min at 4 °C using a SW28 rotor (Beckman). The viral pellet was re-suspended in PBS with 1% BSA and stored at −80’C. The biological titre of the miRNA containing virus was determined by transducing HeLa cells with 10^−2^, 10^−3^ and 10^−4^ dilutions of the vector. 72 h post-transduction, the percentage of GFP positive cells was measured with a Fluorescent-Activated cell sorter (FACS, LSRII). For the repeat containing virus, HeLa cells where transduced with 10^−2^, 10^−3^ and 10^−4^ dilution of the vector. 72 h post-transduction, cells were fixed and immunofluorescence stained with V5 antibody as detailed later to obtain a biological titre. The biological titre is expressed as the number of transducing units per ml (TU/ml) and is calculated as follows: Vector titre = [(% positive cells × total number of cells) × dilution factor × 2].

#### *D. melanogaster* stocks and husbandry

Flies were raised under standard conditions in a humidified, temperature-controlled incubator with a 12 h:12 h light:dark cycle at 25 °C, on standard food consisting of agar, cornmeal, molasses, propionic acid and yeast. Transgene expression was induced using the ubiquitous da-GAL4 driver or tissue specific nSyb-GAL4 or D42-GAL4. The following strains were obtained from the Bloomington D. melanogaster Stock Center (RRID:SCR_006457): da-GAL4 (RRID:BDSC_55850), nSyb-GAL4 (RRID:BDSC_51635), D42-GAL4 (RRID:BDSC_8816). UAS-G4C2×3 and UAS-G4C2×36 lines were a gift from Adrian M. Issacs (UCL Queen Square Institute of Neurology) (*16*). For treatment with CPP, crosses were set up in standard food containing 10 μM Ctrl, SRSF1 WT- or m4-CPPs (Fig. 4).

#### Locomotor assays

The startle induced negative geotaxis (climbing) assay was performed using a counter-current apparatus. Briefly, twenty 2-day-old male adult flies treated or not with 10 μM Ctrl, WT and m4 CPPs were placed into the first chamber, tapped to the bottom, and given 10 s to climb a 19 cm distance. The weighted performance of several group of flies for each genotype was normalized to the maximum possible score and expressed as Climbing index (*28*). For larval crawling assays, wandering third instar larvae treated or not with 10 μM Ctrl-, WT- and m4-CPPs were placed in the centre of a 1% agar plate and left to acclimatise for 30 seconds, after which the number of peristaltic waves that occurred in the following minute were recorded.

#### Recombinant protein purification and GST Pull-down assay

All recombinant proteins were expressed in *E.coli* BL21 (DE3)-RP cells and induced by isopropyl – β-D-thiogalactoside (IPTG, Sigma). Recombinant GB1-6His-SRSF1 (11-196) was purified by IMAC chromatography on TALON/Cobalt beads (Clontech) with the following buffers (Lysis Buffer: 50 mM Tris pH8.0, 1M NaCl, 0.5% Triton X-100; Wash Buffer: 50 mM Tris pH8.0, 1M NaCl, 5 mM Imidazole; Elution Buffer: 20 mM Tris pH8.0, 0.5 M NaCl, 0.2 M Imidazole). Protein concentration was determined by Bradford protein assay (*BioRad*). For pull-down reaction, 0.25 g IPTG-induced GST-NXF1:p15 bacteria pellets was lysed by sonication in phosphate buffered saline (PBS) with 0.5% Triton X-100 and supplemented with protease inhibitor (Complete™, EDTA-free Protease Inhibitor Cocktail, *Roche*) and 2 mM phenylmethanesulfonyl fluoride (PMSF, Sigma), and immobilized on GST Sepharose 4B beads (GE Healthcare). For cell permeable peptide binding reactions, different concentrations of CPPs were incubated with immobilized GST-NXF1:p15 in 250 μl of PBS with 0.5 % Triton X-100 and RNase A (0.04 mg/ml) at 4°C for 1 hr. For competition assay, 1.2 μg of purified SRSF1 (11-196)-6His was incubated with immobilized GST-NXF1:p15 in 250 μl of PBS containing 0.5 % Triton X-100 and RNase A (0.04 mg/ml) and different concentrations of CPPs at 4 °C for 1 hr. The GST-NXF1:p15-interacting proteins were eluted with GSH elution buffer (50 mM Tris, 100 mM NaCl, 40 mM reduced glutathione, pH 7.5) prior to SDS-PAGE electrophoresis and analysed by Coomassie blue staining and immunoblotting.

#### Isothermal Titration Calorimetry (ITC) measurements

The nucleotide sequence encoding NXF1 amino-acids 1-198 was codon optimized for expression in *E. coli,* custom synthesized by GenScript and cloned into pET-28a(+)-TEV vector (Cloning site: NheI/XhoI) containing N-terminal hexahistidine (6His) tag followed by a tobacco etch virus (TEV) protease cleavage site. Cloned 6His-NXF1 1-198 was transformed into *E. coli* BL21(DE3) pLysY/Iq (New England Biolabs) cells and subsequent colonies were used for the initial culture growth at 37 °C in TY media supplemented with Kanamycin antibiotic resistance (50 μg/ml). Bacteria from overnight culture were used to seed 1 l culture for natural abundance of protein. Expressed cells were harvested (4000 x g) and lysed into lysis buffer consisting of 50 mM Tris (pH 7.5), 500 mM NaCl, 5 mM imidazole and 5% (v/v) glycerol and frozen at −80° C in the presence of added lysozyme. Frozen cells were lysed by sonication and soluble protein fractions were separated by ultracentrifugation (20000 x g) for 30 min at 4 °C. The fractions were filtered through 0.45 μM syringe filter (MercMillipore Inc) to remove supernatant and cellular debris. The filtered supernatant was loaded onto 2 ml Ni-NTA Agarose affinity resin (Qiagen UK). The resin than washed with 20 column volume lysis buffer and 10 column volume of same buffer but with 25 mM of imidazole. The proteins were eluted by buffer consisting of 50 mM Tris (pH 7.5), 500 mM NaCl, 250 mM imidazole and 5% (v/v) glycerol. The protein fractions were pooled and buffer exchanged to 20 mM sodium phosphate (pH 6.5), 300 mM NaCl using PD10 desalting column (GE Healthcare). Protein purity was checked by NuPAGE Bis-Tris (Invitrogen) and concentration was determined by absorbance at 280 nm with extinction coefficients obtained using ProtParam tool (http://web.expasy.org/protparam). The protein was concentrated using 3000 MWCO Amicon centrifugal filter units (MerckMillipore Inc). Synthetic FAM-labelled SRSF1 peptides (purity >99%) were purchased from *ThermoFisher Scientific UK*.

For Isothermal titration calorimetry (ITC), protein samples were thoroughly dialyzed overnight into 20 mM sodium phosphate (pH 6.5), 300 mM NaCl. ITC measurement was performed on iTC 200 microcalorimeter (Malvern Instrument) at 298K. The titration cell was filled with 25 μM NXF1 1-198 and syringe was filled with 1 mM peptide (Table S1). After the initial delay of 360 sec a first injection of 0.4 μl followed by 3 μl injection of 13 titrations of peptides with NXF1(1-198) were carried from rotating syringe at a speed of 750 rpm to titrant cell containing NXF1(1-198). The delay between each injection was 180 sec and ITC data was recorded with high sensitivity mode. The data was analysed using the *Origin* software.

#### MTT cell proliferation assay

HEK cells were split into 24-well plates (25,000 cells per well). Each plate contained 4 wells with only media to serve as a blank and 4 wells/treatment. Cells were transfected with either 500 ng pcDNA3.1/G4C2_45_-3xV5 or C4G2_43_-3xV5. 6 h post transfection, media was replaced and 0, 0.25, 0.5 or 1 μM of Ctrl-, m4- or WT-CPPs added. 72 h post transfection, 250 mg Thiazolyl Blue Tetrazolum Bromide reagent (MTT) was added to each well and incubated in the dark at 37 °C for 1 h. Cells were subsequently lysed with equal volume MTT lysis buffer (20%SDS, 50% dimethylformamide (DMF)) and incubated in the dark, shaking at room temperature for 1 h. Absorbance at 595 nm was assessed with a PHERAstar FS (BMG Labtech). Absorbance data was retrieved using PHERAstar MARS (BMG Labtech).

#### Nuclear/Cytoplasmic fractionation for qRT-PCR and western immunoblotting from HEK293T cells and patient-derived iNeurons)

HEK cells were split into 6-well plates (200,000 cells per well) and transfected for 72 h with either 2 μg pcDNA3.1/G4C2_45_-3xV5 or C4G2_43_-3xV5. 6 h post transfection, the media was replaced and 0, 0.5 or 1 μM of Ctrl- or WT-CPP was added. For patient-derived neurons, induced neural progenitor cells (iNPCs) were differentiated to iNeurons using a modified version of the protocol (*14*) as previously described (*5*). In brief, 100,000 iNPCs were plated in each well of 6-well plates coated with fibronectin (Millipore) and expanded to 70-80% confluence. Once they reached this confluence, iNPC medium was replaced with neuron differentiation medium (DMEM/F-12 with Glutamax supplemented with 1% N2, 2% B27 (Gibco) containing 2.5 μM of DAPT (Tocris) to determine differentiation towards neuronal lineage on day 1. On day 3, the neuron differentiation medium is supplemented with 1 μM retinoic acid (Sigma), 0.5 μM smoothened agonist (SAG) (Millipore) and 2.5 μM forskolin (Sigma) for 7 days. This protocol leads to typical yields of 70% β-III tubulin (Tuj1) positive cells. 5 μM CPPs were added into differentiating iNeurons on day 7 for 72 hours prior to cell harvest. Induced motor neurons (iMNs) cannot be generated in a sufficient number for this assay.

For the cytoplasmic fractionation, 3 wells (HEK cells) or 6 wells (iNeurons) of a 6-well plate were collected in DEPC PBS using a cut tip and pelleted by centrifugation at 400 x *g* for 5 min. Cell pellets were quickly washed with hypotonic lysis buffer (10 mM HEPES pH 7.9, 1.5 mM MgCl2, 10 mM KCl, 0.5 mM DTT) and lysed for 10 min on ice in hypotonic lysis buffer containing 0.16 U μl^−1^ Ribosafe RNase inhibitors (Bioline), 2 mM PMSF (Sigma) and SIGMAFAST Protease Inhibitor Cocktail tablets, EDTA free (Sigma). All lysates underwent differential centrifugation (1,500 *g*, 3 min, 4 °C then 3,500 *g*, 8 min, 4 °C and then 17,000 *g*, 1 min, 4 °C) transferring the supernatants to fresh tubes after each centrifugation. Nuclear pellets obtained after centrifugation at 1,500 *g* for 3 min were lysed in Reporter lysis buffer (Promega) for 10 min on ice before centrifugation at 17,000 *g*, 5 min, 4 °C. Total fractions were collected from 1 well (HEK cells) or 3 wells (iNeurons) of a 6-well plate in Reporter lysis buffer containing 16 U μl^−1^ Ribosafe RNase inhibitors (Bioline), 2 mM PMSF (Sigma) and protease inhibitors prior to lysis for 10 min on ice before centrifugation at 17,000 *g*, 5 min, 4 °C. Equal volumes of total, nuclear and cytoplasmic lysates were subjected to western immunoblotting using SSRP1 and HSPA14 antibodies (HEK293T cells) or SSRP1 and Anti-Beta-Tubulin III/TUJ1 antibodies (iNeurons).

Total and fractionated extracts were added to PureZOL™ (BioRAD) to extract the RNA. Briefly, total and nuclear lysates was cleared by centrifugation for 10 min at 12,000 *g* at 4°C. One fifth the volume of chloroform was added and tubes were vigorously shaken for 15 sec. After 10 min incubation at room temperature, tubes were centrifuged 12,000 *g*, 10 min, 4°C and supernatants collected. RNA was precipitated overnight at −20°C with equal volume isopropanol and 1 μl glycogen (5 μg μl^−1^, Ambion) and subsequently pelleted at 12,000 *g,* 20 min, 4°C. Pellets were washed with 70% DEPC ethanol and re-suspended in DEPC water. All PureZol™ extracted RNA samples were treated with DNaseI (Roche) and quantified using a Nanodrop (Nano Drop Technologies).

Following quantification, 2 μg RNA was converted to cDNA using BioScript Reverse Transcriptase (Bioline). Human U1 (Fwd: 5’-CCATGATCACGAAGGTGGTT-3’ / Rev: 5’-ATGCAGTCGAGTTTCCCACA-3’) and C9RAN (Fwd 5’-GGGCCCTTCGAACCCCCGTC-3’/ Rev: 5’GGGAGGGGCAAACAACAGAT-3’) qRT-PCR primers used in this study are described in (*6*). qRT-PCR reactions were performed in duplicate using the Brilliant III Ultra-Fast SYBR Green QPCR Master Mix (Agilent Technologies) on a C1000 Touch™ thermos Cycler using the CFX96™ Real-Time System (BioRAD) using an initial denaturation step, 45 cycles of amplification (95 °C for 30 s; 60 °C for 30 s; 72 °C for 1 min) prior to recording melting curves. qRT-PCR data was analysed using CFX Manager™ software (Version 3.1) (BioRAD) and quantified with quantification was performed using the comparative C_T_ method (*29*) and GraphPad Prism (Version 7).

#### Nuclear/Cytoplasmic fractionation for qRT-PCR and western immunoblotting from *D. melanogaster* samples

For the cytoplasmic fractionation, fifteen third instar larvae were homogenised in 400 μl of hypotonic lysis buffer (10 mM HEPES pH 7.9, 1.5 mM MgCl_2_, 10 mM KCl, 0.5 mM DTT) containing 1 μl/ml RNase OUT recombinant ribonuclease inhibitor (Thermo Fisher Scientific, RRID:SCR_008452), 2 mM PMSF (<u>Sigma-Aldrich</u>, RRID:SCR_008988) and complete protease inhibitor cocktail tablets, EDTA free (Sigma-Aldrich, RRID:SCR_008988).

All lysates underwent differential centrifugation (1,000 rpm, 2 min, 4 °C, then 4,000 r.p.m., 3 min, 4 °C and then 13,300 r.p.m., 1 min, 4 °C) transferring the supernatants to fresh tubes after each centrifugation. Nuclear pellets obtained after centrifugation at 4,000 r.p.m. for 3 min were lysed in reporter lysis buffer (Promega, RRID:SCR_006724) for 10 min on ice before drawing up/down with a 25G needle four times. Total fractions were isolated from five third instar larvae in RIPA lysis buffer [50 mM Tris·HCl, 150 mM NaCl, 1 mM EDTA, 1% Triton X-100, 0.5% SDS] with 1 mM PMSF and protease inhibitor mixture (Roche, RRID:SCR_001326). Equal volumes of total, nuclear and cytoplasmic lysates were subjected to western immunoblotting using histone H3 (1:5000, Abcam, RRID:AB_302613) and tubulin (1:5000, Sigma-Aldrich Cat# T9026, RRID:AB_477593) antibodies detected using HRP-conjugated rabbit (1:5000, Invitrogen, RRID:AB_2536530) and mouse (1:5000, Abcam, RRID:AB_955439) secondary antibody respectively.

Total and fractionated extracts were added to TRI Reagent LS (Sigma-Aldrich, RRID:SCR_008988) to extract the RNA. Briefly, total and cytoplasmic lysates was cleared by centrifugation for 10 min at 12,000 *g* at 4 °C. One fifth the volume of chloroform was added and tubes were vigorously shaken for 15 sec. After 10 min incubation at room temperature, tubes were centrifuged 12,000 *g*, 10 min, 4 °C and supernatants collected. RNA was precipitated for 30 min at room temperature with equal volume isopropanol and subsequently pelleted at 12,000 *g,* 20 min, 4 °C. Pellets were washed with 70% DEPC ethanol and re-suspended in DEPC water. All extracted RNA samples were treated with Turbo DNA-free kit (Thermo Fisher Scientific, RRID:SCR_008452) and quantified using a Nanodrop (NanoDropTechnologies).

Following quantification, 500 ng of RNA was converted to cDNA using Maxima H Minus cDNA Synthesis Master Mix (Thermo Fisher Scientific, RRID:SCR_008452) following manufacturer’s instructions. *D. melanogaster* G4C2×36 3’UTR (Fwd: 5’-TTCCAACCTATGGAACTGATGA-3’/Rev: 5’-GGTTTTCCTCATTAAAGGCATTC) and Tub84b (Fwd: 5’-CTTCCTCATCTTCCACTCGTTC-3’/Rev: 5’-ACTCCAGCTTGGACTTCTTG-3’) qRT-PCR primers were used in this study. Quantitative realtime PCR (qRT-PCR) was performed on a CFX96 Touch Real-Time PCR Detection System (Bio-Rad Laboratories, RRID:SCR_008426) and the relative transcript levels of each target gene were normalized against *Tub84b* mRNA levels; quantification was performed using the comparative C_T_ method (*28*), and GraphPad Prism software 7 (GraphPad Prism, RRID:SCR_002798).

#### Co-cultures of patient-derived astrocytes and motor neurons

##### Differentiation of iMotor Neurons (iMNs)

Human patient and control-derived neurons (iNeurons) were differentiated from induced neural progenitor cells (iNPCs) using a modified version of protocol (*14*) as previously described (*5*). In brief, 100,000 iNPCs were plated in a 6-well plate coated with fibronectin (Millipore) and expanded to 70-80% confluence. Once they reached this confluence, iNPC medium was replaced with neuron differentiation medium (DMEM/F-12 with glutamax supplemented with 1% N2, 2% B27 (Gibco) containing 2.5 μM of DAPT (Tocris) to determine differentiation towards neuronal lineage on day 1. On day 3, the neuron differentiation medium is supplemented with 1 μM retinoic acid (Sigma), 0.5 μM smoothened agonist (SAG) (Millipore) and 2.5 μM forskolin (Sigma) for 7 days until Day 10. This protocol leads to typical yields of 70% β-III tubulin (Tuj 1) positive cells. To obtain iMotor Neurons (iMN), ~ 5,000 iNeurons per well were re-plated on 96-well plate coated fibronectin and maintained in iNeuron differentiation medium (containing retinoic acid, SAG and forskolin) supplemented with BDNF, CNTF and GDNF (all at 20 ng/ml) for the last 14 days of differentiation.

##### Differentiation of iAstrocytes

Human patient-derived astrocytes (iAstrocytes) were differentiated from iNPCs as previously described (*5, 14*) and cultured in DMEM glutamax (*Gibco*) with 10% FBS (*Sigma*) and 0.02% N2 (*Invitrogen*) for 5 days. Cells were maintained in a 37°C incubator with 5% CO_2_.

##### Co-cultures of patient-derived iMNs and iAstrocytes

iAstrocytes were lifted at day 5 of differentiation and ~5,000 iAstrocytes were re-plated on iMNs at day 20 of differentiation. Cocultured iMNs and iAstrocyte were maintained in neuron differentiation medium with BDNF, GDNF and CTNF (all at 20 ng/ml) for 4 days. 12 hours after the start of co-cultures (on day 21), 1 or 10 μM CPPs was added to the medium and iMNs/ iAstrocytes were imaged for 72 h at day 22, 23, 24. For SRSF1 knockdown, iMNs and iAstrocytes were separately transduced 48h prior to co-cultures with lentivirus (LV) expressing control or SRSF1-RNAi co-expressing GFP (*6*) at an MOI of 5 at day 18 of iMN differentiation and at day 3 of iAstrocyte differentiation.

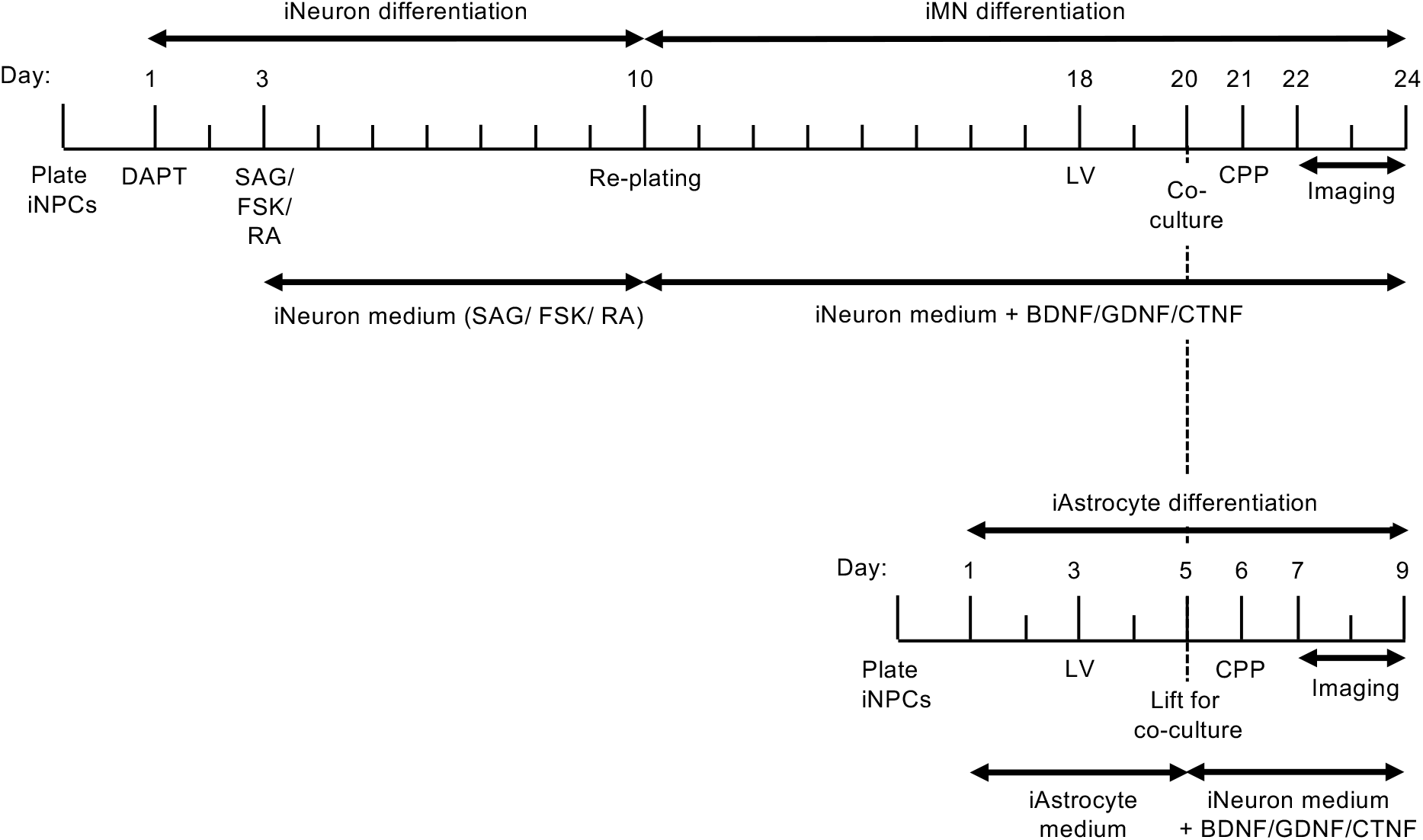

#### Western blot and dot blot analysis

HEK cells cultured in a 24 well plate (50,000 cells/well) were transfected with 700 ng pcDNA3.1/G4C2_45_-3xV5 or C4G2_43_-3xV5 using 3.5 μg PEI/ml media and one tenth medium volume OptiMEM. 6 h post transfection, media was replaced and 0, 0.1, 0.25, 0.5, 0.75, 1 or 2.5 μM Ctrl-, m4- or WT-CPPs were added. Proteins were extracted 72 h post-transfection. Cells were washed in ice-cold phosphate-buffered saline (PBS) and scraped into ice-cold lysis buffer (50 mM Hepes pH7.5, 150 mM NaCl, 10% glycerol, 0.5% Triton X-100, 1 mM EDTA, 1 mM DTT, protease inhibitor cocktail (Sigma)). Cells were lysed on ice for 10 min followed by centrifugation at 17,000 *g* at 4 °C for 5 min. Protein extracts were quantified using Bradford Reagent (BioRAD), resolved by SDS–PAGE, electroblotted onto nitrocellulose membrane and probed using the relevant primary antibody.

Cultured cortical neurons were transduced with LV-G4C2_45_-3xV5 or LV-C4G2_43_-3xV5 and either 5 MOI LV-Ctrl RNAi/ LV-SRSF1-RNAi or 1 μM/ 5 μM Ctrl-, m4- or WT-CPPs. 16 h post transduction, media was replaced and CPPs added again to appropriate samples. Proteins were extracted 72 h post-transfection. iAstrocytes were cultured in media containing Ctrl, m4- or WT-CPPs for 72 h. Cortical neurons and iAstrocytes were washed in ice-cold PBS and scraped into ice-cold lysis as detailed above. Cells were passed through a 25G needle 10 times to improve lysis efficiency and left to lyse on ice for 10 min followed by centrifugation at 17,000 *g* at 4 °C for 5 min. Protein extracts for cortical neurons were quantified using Bradford Reagent, resolved by SDS-PAGE, electroblotted onto nitrocellulose membrane and probed using the relevant primary antibody. Protein extracts from iAstrocytes were subjected to dot blot analysis. Extracts were quantified using Bradford Reagent and 50 μg total protein extracts prepared in ice-cold lysis buffer were loaded onto a nitrocellulose membrane using a microfiltration apparatus (Biorad), sliced into strips and analysed by immunoblotting.

*D. melanogaster* third instar larvae bearing 3 or 36 G4C2 repeats fed with either Ctrl, m4- or WT-CPP were frozen in liquid nitrogen and then lysed in ice-cold lysis buffer detailed above using Eppendorf micropestles. Lysed larvae were incubated on ice for 10 min followed by centrifugation at 17.000 *g* for 10 min at 4 °C. Protein extracts were quantified using Bradford Reagent and subjected to dot blot analysis as detailed above.

For all studies, poly-Gly-Pro [1:2000 dilution] (Proteintech #24494-1-AP), poly-Gly-Arg [1:2000] (Proteintech #23978-1-AP), poly-Pro-Arg [1:2000] (Proteintech #23979-1-AP), SRSF1 [1:1000] (Cell signalling #14902), HSPA14 [1:2000] (Abcam #ab108612), Histone H3 [1:5000] (Abcam #ab1791) and V5 clone D3H8Q [1:500] (Cell Signalling #13202S) primary antibodies were detected with horseradish peroxidase (HRP)-conjugated rabbit secondary antibody (Promega), whilst α-tubulin (Insight, clone DM1A #Sc32293, [1:2000] for cell work / Sigma-Aldrich, clone DM1A #T9026 for *Drosophila* work), poly-Gly-Ala [1:500] (kindly provided from Prof Dieter Edbauer), V5 [1:5000] (Invitrogen #R96025), NXF1 [1:2000] (Abcam #ab50609), SSRP1 [1:500] (Abcam #ab26212), FLAG [1:2000 dilution] (Sigma F1804, clone M2) and actin [1:5000] (Abcam #ab8224) antibodies were detected using HRP-conjugated mouse secondary antibody (Promega). Poly-Pro-Ala [1:1000] (Merck clone 14E2 #MABN1790) was detected using HRP-conjugated rat secondary antibody (Promega). Anti-Beta-Tubulin III [1:1000] (Merck #AB9354) was detected using HRP-conjugated chicken secondary antibody (Promega). Uncropped western blot images can be provided on request.

#### Immunofluorescence

For peptide uptake immunofluorescence assays, HEK293T cells (2,000 cells per well) and primary rat cortical neurons (4,500,000 neurons per 24-well plate) were cultured in black optical 96 well plates. Ctrl, m4- or WT-CPP at concentrations of 0, 1, 5 or 10 μM were added to the media 24 h after plating. Immunofluorescence staining of HEK and rat cortical neurons was performed 48 h after CPP addition as described previously (6, *25*) with the exception that cells were blocked with 4% goat serum in PBS for 2 h at room temperature and incubated overnight at 4 °C with the V5 antibody [1:1,000 dilution] (ThermoFisher Scientific #R96025) in PBS containing 4% goat serum. Cells were washed three times with PBS containing 4% goat serum and incubated for 1 h with Beta III tubulin antibody [1:1000 dilution] (Merck #9354) in PBS containing 4% goat serum. Cells were then washed three times with PBS containing 4% goat serum and incubated for 1 h in 4% goat serum with goat anti-mouse secondary antibody [Alexa Fluor 488 (1:1,000 dilution); ThermoFisher Scientific] to detect V5 and anti-chicken secondary antibody [Alexa Flour 647 (1:1000 dilution); ThermoFisher Scientific] to detect TUJ1. Cells were subsequently stained with Hoechst 33342 for 10 min at room temperature, washed 3 times with PBS and imaged using the Opera Phenix high content screening system (Perkin Elmer). For any well a minimum of 9 random fields using the 20x objective were captured in 3 focal planes, meaning an excess of 3000 cells in total were obtained for any well to calculate percent uptake. Quantification was assessed using the Columbus Image data storage and analysis system (Perkin Elmer) software, nuclei were identified using Hoechst (blue channel), and cytoplasm was masked using beta III tubulin/TUJ1 (red channel) and peptide uptake was measured (green channel) in both nucleus and cytoplasm. For representative images of cellular distribution, cells were captured using 40x objective.

For DPR and peptide immunofluorescence, primary rat cortical neurons were isolated and cultured in 24-well plates with coverslips (4,500,000 cells per plate). Cells were transduced with LV-G4C2_45_-3xV5 or LV-C4G2_43_-3xV5 and either 5 MOI LV-ctrl RNAi/LV-SRSF1-RNAi or 5 μM Ctrl or WT-CPP. 16 h post transduction, the media was replaced and control or SRSF1 WT CPP added to appropriate samples. Immunofluorescence of cortical neurons was performed 7 days after transduction as described previously (*5*). Cells were imaged using a Zeiss Axioplan 2 microscope. The experiments were repeated three times and within each experiment, each condition was examined with >10 fields of view with a minimum of 100 cells. All analysis was performed blinded to experimental condition to test whether SRSF1 depletion led to reduced DPR synthesis.

#### Co-immunoprecipitation

Cells were split into two 10 cm plates/treatment (1.5 × 10^6^ cells per plate) and transfected with 15 μg p3xFLAG or p3xFLAG/NXF1 using 3 μg PEI/1 μg DNA and one tenth medium volume OptiMEM. After transfection 1μM control / SRSF1 CPP was added to the media. Proteins were extracted from HEK cells 48 h post-transfection. Cells were washed in ice cold PBS, scraped into 500 μl ice cold lysis buffer, passed through a 2**5**G gauge needle 10 times and left to lyse on ice for 10 min. Lysed cells were cleared by centrifugation at 17,000 *g* at 4 °C for 5 min and protein extracts were quantified using Bradford Reagent. 2 mg total protein in 1 ml lysis buffer was incubated with 20 μl Anti-FLAG^®^ M2 Magnetic Beads (Sigma M8823) (which had been blocked overnight with 1% BSA in IP lysis buffer) for 2 h at 4 °C on a rotating wheel. Beads were washed 5 times with lysis buffer and eluted in 50 μl IP lysis buffer supplemented with 100 μg ml^−1^ 3xFLAG peptide (Sigma #F4799) for 30 min at 4 °C on a rotating wheel. 30 μg total protein and 15 μl eluates were subjected to western immunoblotting using FLAG, SRSF1, V5 and α-tubulin antibodies.

#### Statistical analysis of data

One-way and two-way ANOVA (analysis of variance) with Tukey’s correction for multiple comparisons were used for most experiments with the following exceptions: DPR analysis in primary neurons used Fisher’s exact test; *D. melanogaster* climbing assay was analysed by Kruskal-Wallis non-parametric test with Dunn’s correction for multiple comparisons; For *D. melanogaster* crawling assays One-way ANOVA with Bonferroni’s multiple comparation test was used; G4C2×36 transcripts in *D. melanogaster* were quantified using One-way ANOVA with Sidak’s correction for multiple comparison test. No randomization was used in the animal studies. Data were plotted using GraphPad Prism 7. Significance is indicated as follows; NS: non-significant, p≥0.05; * p<0.05; ** p<0.01; *** p<0.001; **** p<0.0001. DPR-positive neurons, crawling and climbing assays were analysed in a blinded manner. Several researchers were involved in producing replicate experiments for qRT-PCR and western blot data.

## Supplementary Figures

**Fig. S1.**
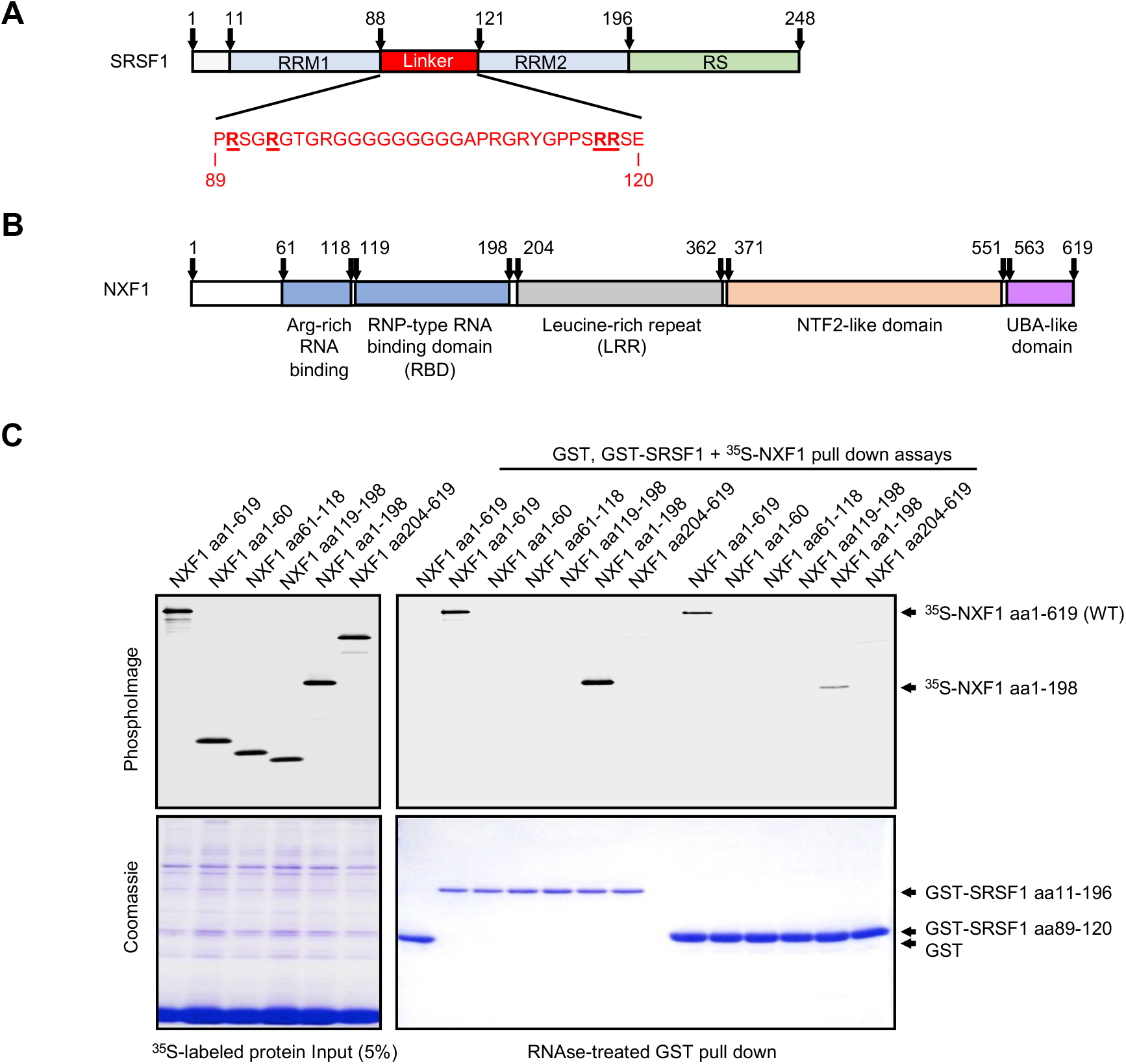
The unstructured linker region of SRSF1 comprising amino-acids 89-120 of SRSF1 interacts with the RNA binding domain of NXF1. (**A**) Schematic diagram of SRSF1 highlighting the RNA recognition motifs (RRM), the NXF1-interacting linker region and the RS domain which is rich in serine and arginine residues. N **timbe**rs depict amino-acid (aa) position. (**B**) Schematic diagram of NXF1 highlighting the RNA binding regions comprising an arginine-rich RNA-binding domain and the RNP-type RNA binding domain (aa1-198) (*12*), a leucine-rich repeat domain (aa 204-362), a NTF2-like domain (aa 371-551) which heterodimerizes with p15/NXT1 (*25, 26*) and a UBA-like domain (aa 563-619) which associates wich nucleoporins together with the NTF2-like scaffold (*30*). The p15 protein heterodimerizes and stabilises NXF1 to stimulate the NXF a-depandent RNA nuclear exp ort (*26*). Ia will therefore be co-expressed with NXF1 in the *in vitro* GST-NXF1 pull down assays performed in this study. (**C**) Recombinant bacteriallydexpressed GST and GST-NXF1:p15 pull down assays were performed in presence of RNase and mammdlian recombinant ^35^S-labelled NXF1 protein full leng(_d_ (WT) or various domains expressed in rabbit reticulocytes.

**Fig. S2.**
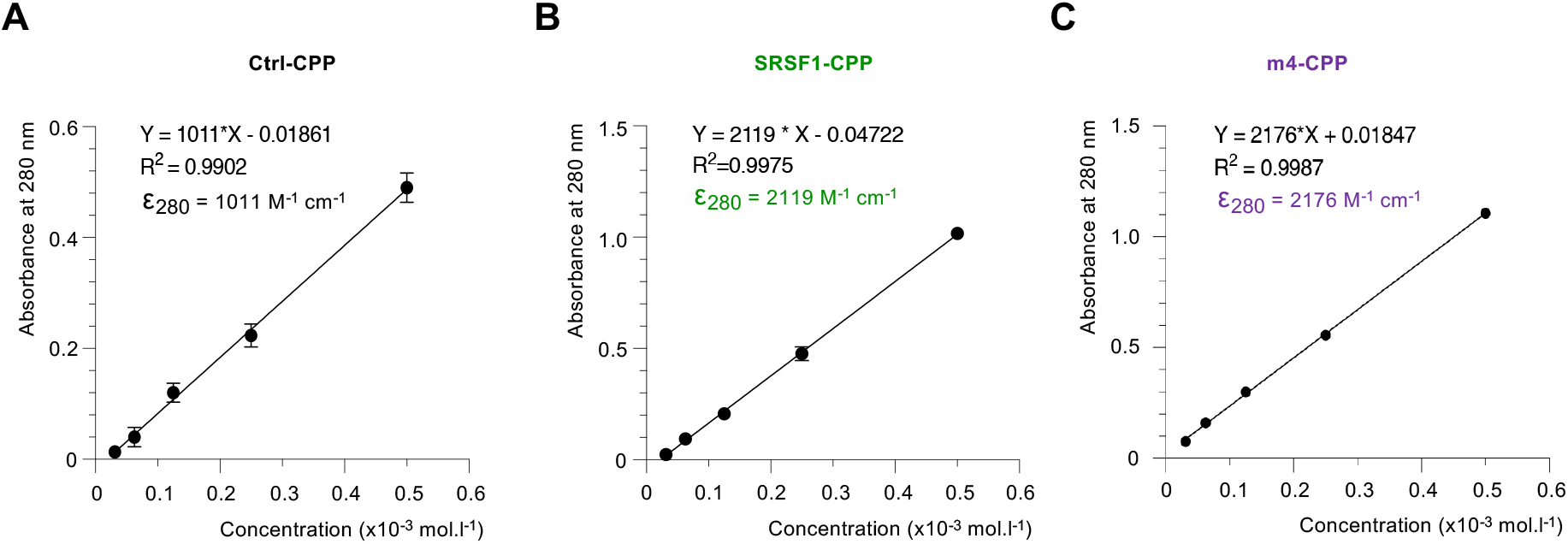
Measuring the concentrations of cell permeable peptides. The cell permeable peptides were custom synthesized by *ThermoFisher Scientifc* at >90% purity and resuspended in PBS at 1 mM. Absorbance of 1:2 serial dilutions of the CPPs (0.5 to 0.03125 mM) was measured at 280 nm in triplicate. According to the Beer-Lambert Law A = ε x l x C (A: Absorbance, ε: molar extinction coefficient, l: optical path, C: concentration), the slope of the graph A = f(C) corresponds to ε x 1 where l = 1 cm. Error bars represent standard deviation. The experimental molar extinction coefficients were determined in PBS for Ctrl-CPP (**A**, ε_280_ = 1011 M^−1^ cm^−1^), WT-CPP (**B**, ε_280_ = 2119 M^−1^ cm^−1^) and m4-CPP ((3, ε_280_ = 2176 M^−1^ cm^−1^). In comparison, the theoreticar vulue computed in water (https://web.expasy.org/protparam/) were respectively 1490, 2980 and 2980 M^−1^ cm^−1^. We used the determined experimental molar extinction coefficients and measured the absorbance at 280 nm of sto ck CPP sol utions to accurately quantify their concentrations across batches.

**Fig. S3.**
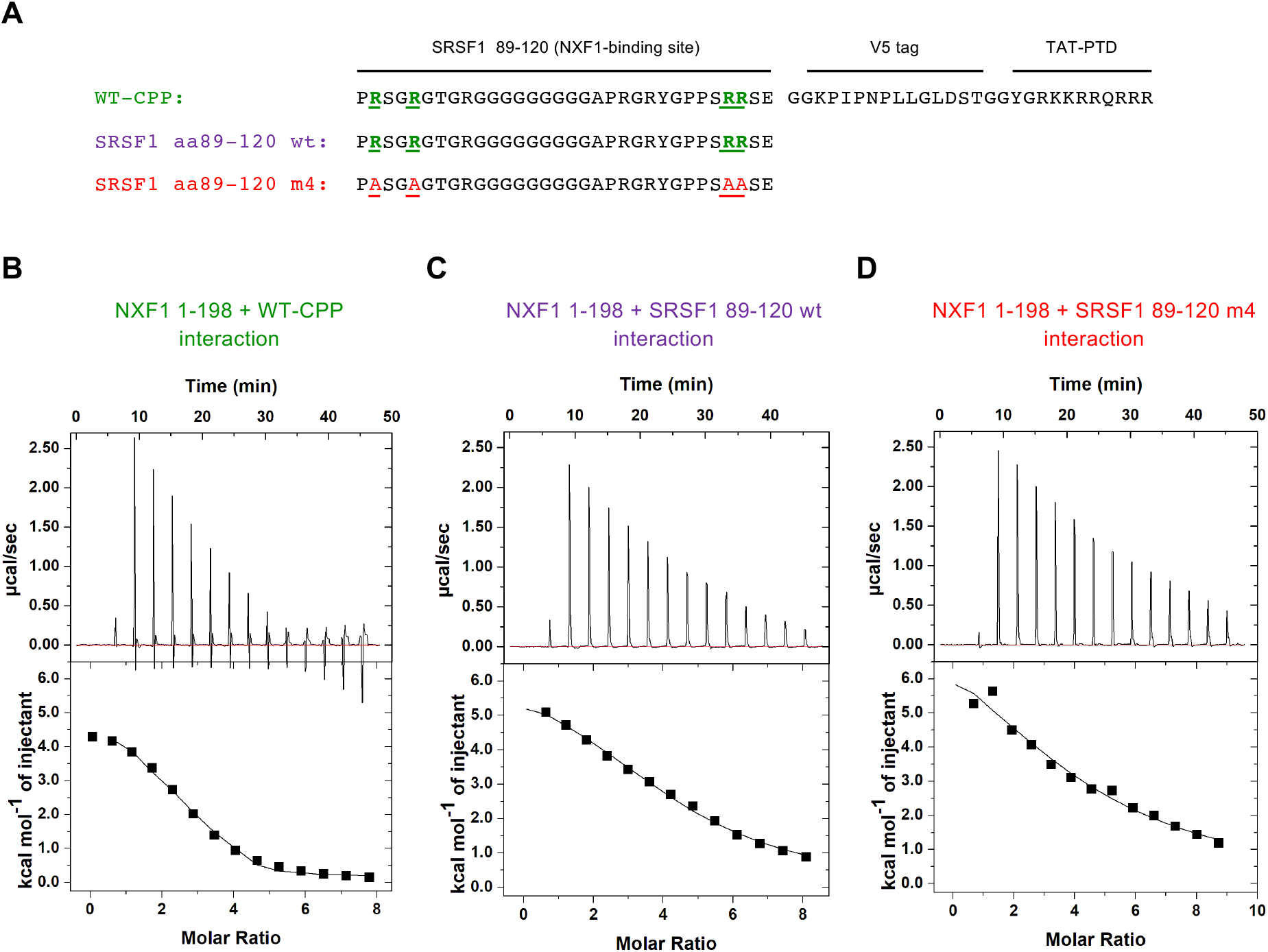
Mutations of SRSF1 arginines 90, 93, 117, 118 (m4) reduce the binding affinity of SRSF1 for NXF1. **(A)** Minimal SRSF1 wild type (wt) and m4 peptide sequences interacting with the SRSF1-binding site of NXF1 (NXF1 aa 1-198). The m4 peptide comprises substitutions of arginine 90, 93, 117, 118 involved in NXF1-binding into alanine (underlined). **(B-D)** Isothermal titration calorimetry of peptides binding to NXF1. Purified recombinant NXF1 aa 1-198, which corresponds to the SRSF1-binding site, was incubated with successive injections of SRSF1 WT-CPP (B) aa 89-120 wt (C) or SRSF1 aa 89-120 m4-mutant (D) peptides. The upper panels show the raw calorimetric data and the bottom panels are; binding isotherms as a function of the peptide:NXF1 m olar ratio.

**Fig. S4.**
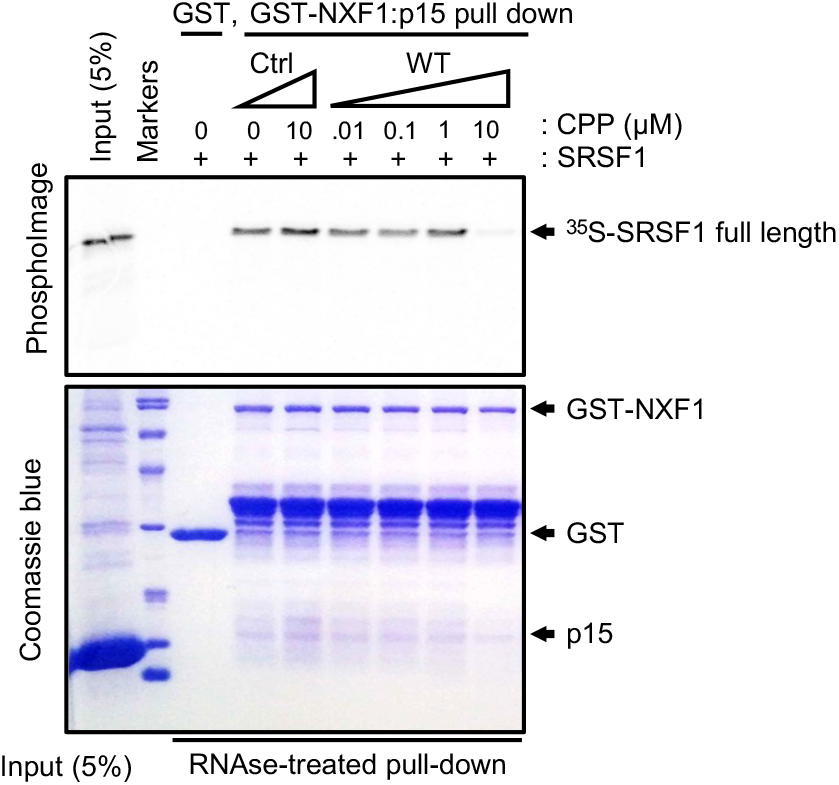
SRSF1 WT-CPP competes with the interaction of SRSF1 full length and NXF1. Recombinant bacterially-expressed GST and GST-NXF1:p15 pull down assays were performed using mammalian recombinant SRSF1 and increasing concentrations of Ctrl- or WT-CPPs.

**Fig. S5.**
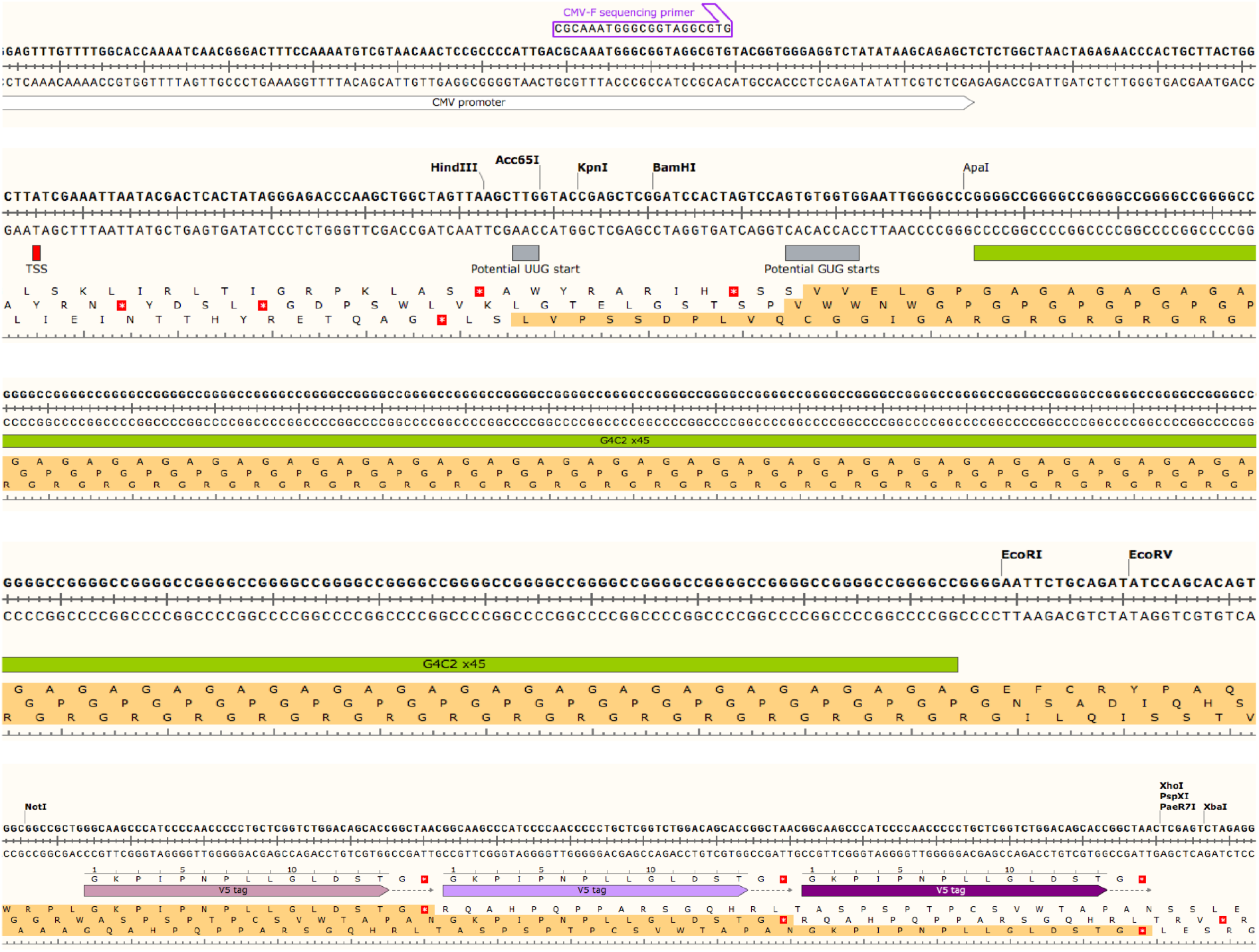
Engineering RAN-dependent G4C2x45 repeat transcripts with 3xV5 tags in all open reading frames. Concatemerized and annealed G4C2/C4G2x15 DNA oligonucleotides blunted with Mung bean nuclease were cloned into the Klenow-filled site EcoRI of pcDNA3.1 to build pcDNA3.1-G4C2x45. A NotI “XbaI cassette encoding 3xV5 tags and 3 stop codons ic all frames was digested from a synthetic custom-synthesized plasmid (*ThermoFisher Scientific*) and subcloned into the NotI/XbaI site of pcDNA3.1-G4C2x45. Stmger sequencing is available on request. Note the ebsence of canonical start cedeiis from the transcription start site (TSS) to investigate RAN-dependent translation of sense *C9ORF72*-repeat transcripts. The one letter amico-acid code sequences highlighted in orange indicate the DPRs respectively produced in the 3 reading frame’s.

**Fig. S6.**
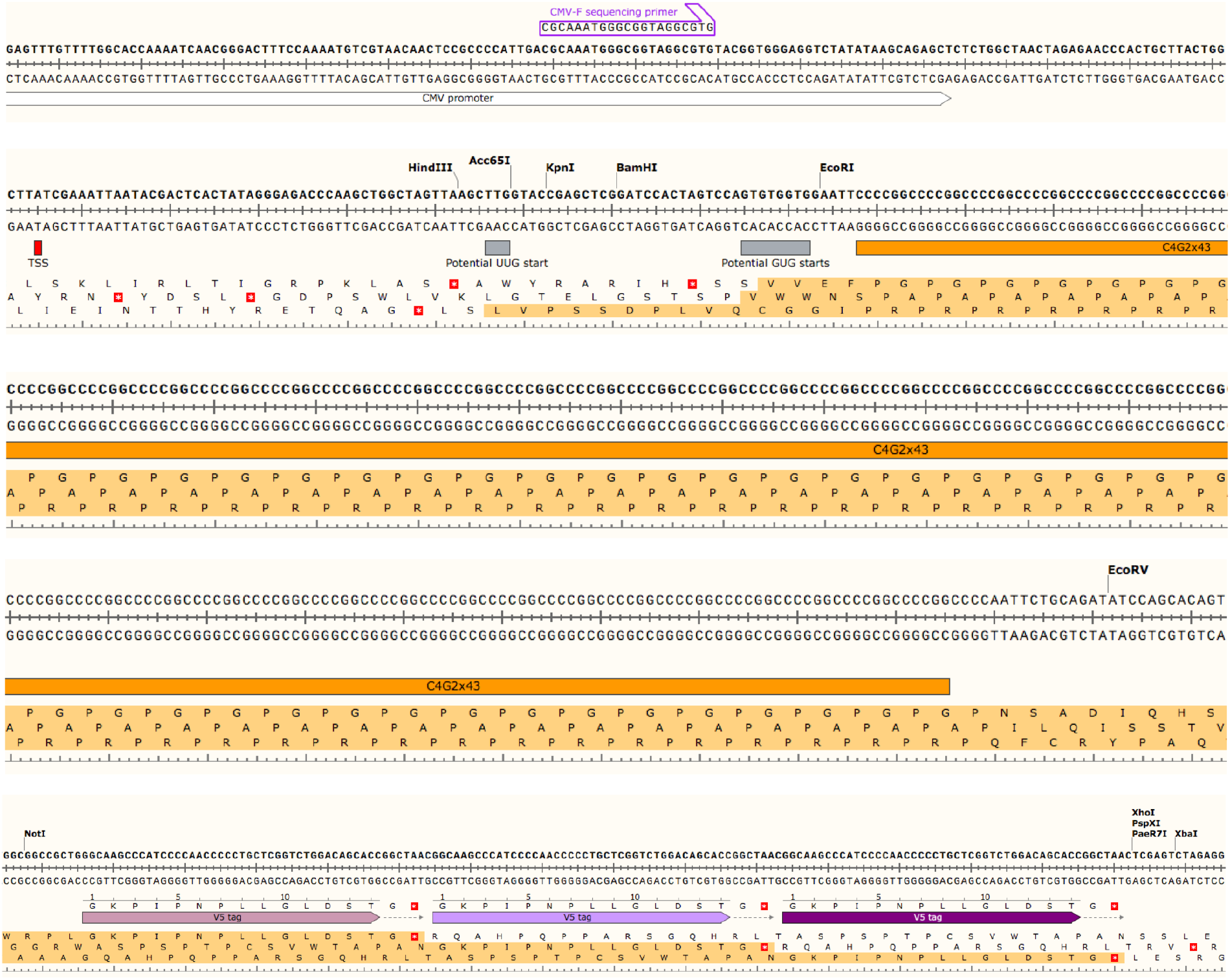
Engineering RAN-dependent C4G2x43 repeat transcripts with 3xV5 tags in all open reading frames. Concatemerized and annealed G4C2/C4 G2x 1 5 DNA oligonucleotides blunted with Mung bean nuclease were cloned into the Klenow-filled site *l*ieoRI of pcDNA3.1 to build pcDNA3.°-C4G2x43. A NotI/XbaI cassette encoding 3xV5 tags nnd 3 stop codons in all frames was digested from a synthetic custom-synthesized plasmid (*ThermoFisher Scientific*) and subcloned into the NotI/XbaI site of pcDNA3.1-C4G2x43. Sanger sequencing is available on request. Note the abeence of canonical AUG start codone from the transcription start site (TSS) to investigate RAN-dependent translation of sense *C9ORF72*-repeat transcripts.The one letter amino-acid code sequences highlighted in orange indicate the DPRs respectively produced in the 3 reading frames.

**Fig. S7.**
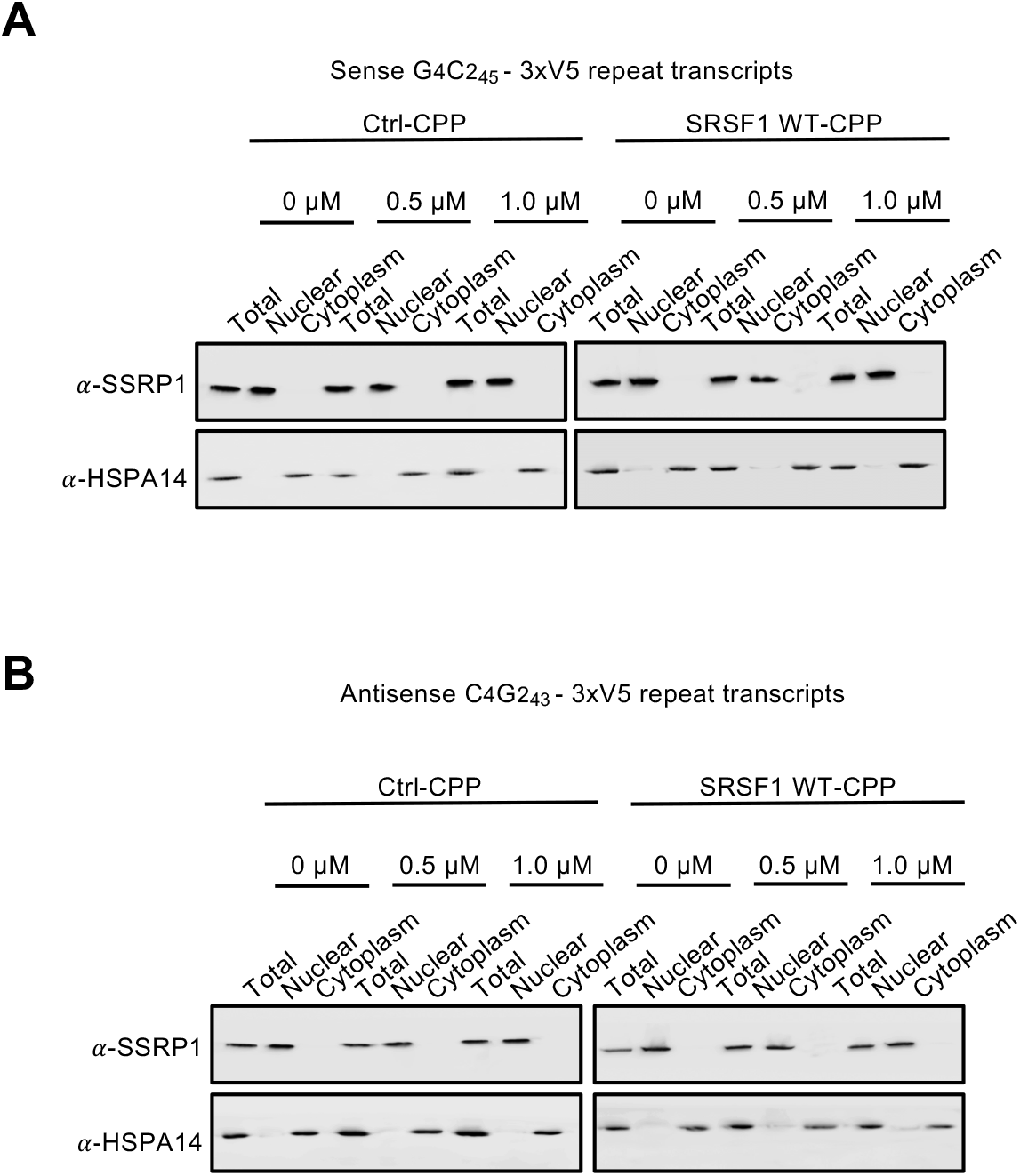
Preparation of total, nuclear and cytoplasmic fractions for quantifying the nuclear export of reporter repeat transcripts in human cell models of C9ORF72-ALS. (**A, B**) Western blots of total, nuclear and cytoplasmic fractions isolated from HEK293T cells transfected for 72h with G4C2_45_-3xV5 (A) or C4G2_39_-3xV5 (B) plasmids and treated with increasing crn centrations of Ctrl ar SR SF1 WT CPPs were probed with nuclear cliromatin-associated SSRP1 factor and predominantly cytoplasmic heat-shock HSPA14 protein.

**Fig. S8.**
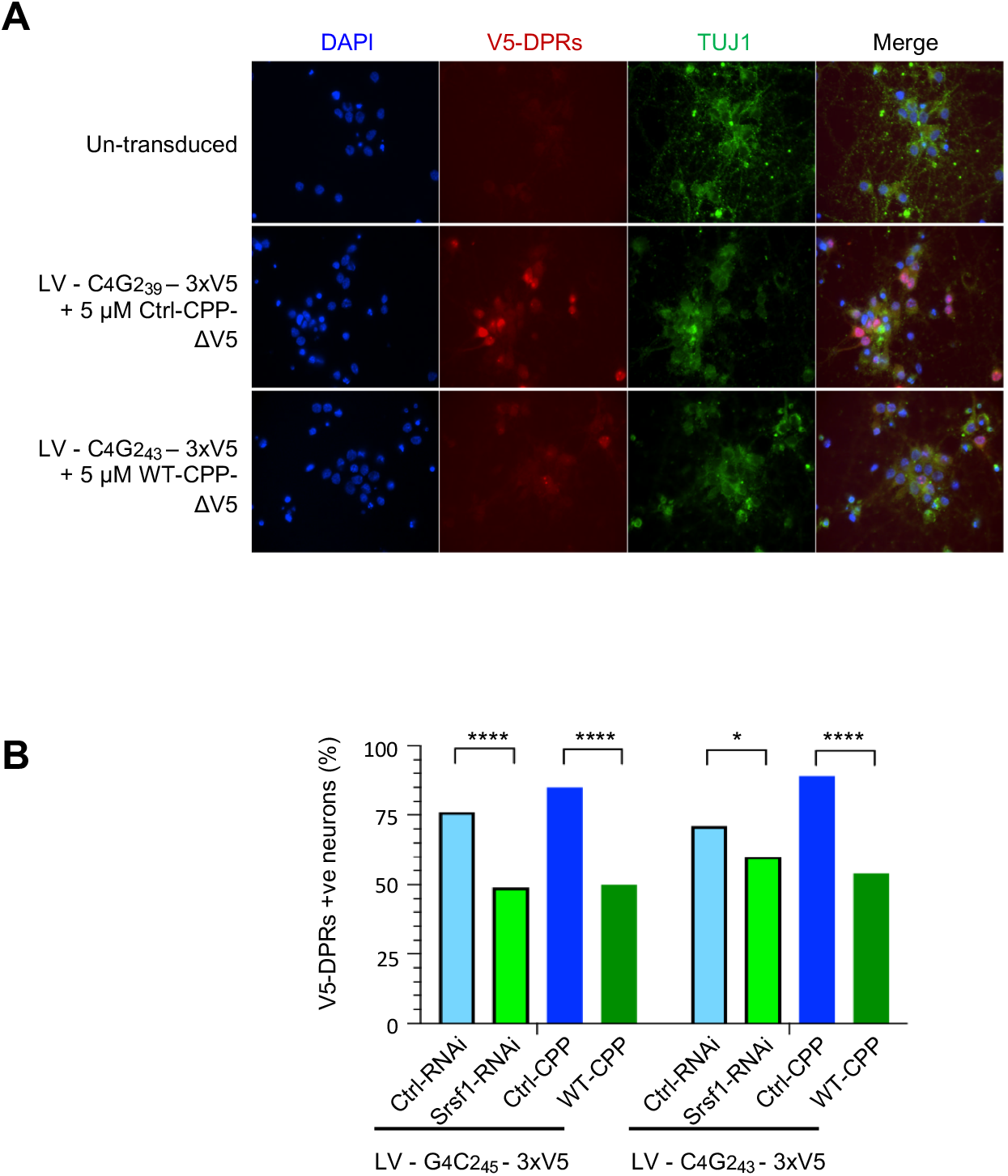
SRSF1 WT-CPP inhibits RAN-translation of DPRs in primary rat cortical neurons. (**A**) Rat cortical primary neurons transduced or not with a lentivirus expres sing C4G2_43_-3xV5 repeat transcripts were treated or not with Ctrl or WT CPPs lacking V5 tags prior to V5 (in the red channel) and TUJ1 (in the green channel) immunofluorescence microscopy. DAPI was used to stain the nuclei. Data are shown for neurons transduced with virus expressing antisense DPRs. (**B**) Sam experiments as in panel D but including lentivirus expressing G4C2_45_-3xV5 repeat transcripts, Ctrl-RNAi or SRSF1-RNAi. For each condition, V5-DPRs positive neurons were counted blinded in at least 10 microscopic fields for at least 100 neurons in biological tripticate experimenta prior to use a Fisher’s exact test to assess statistical significance of either SRSF1 depletion or WT-CPP treatment. Ctrl-RNAi versus SRSF1-RNAi conditions respectively generatad p-values of 2.3E-05 (****) and 0.02 (*) for transductions with sense and antisense repeat transcripts. Ctrl-CPP versus SRSF1 WT-CPP conditions respectively generated p-values of 9.5E-09 (****) and 1.8E-11 (****) for transductions with sense and antisense repeat transcripts.

**Fig. S9.**
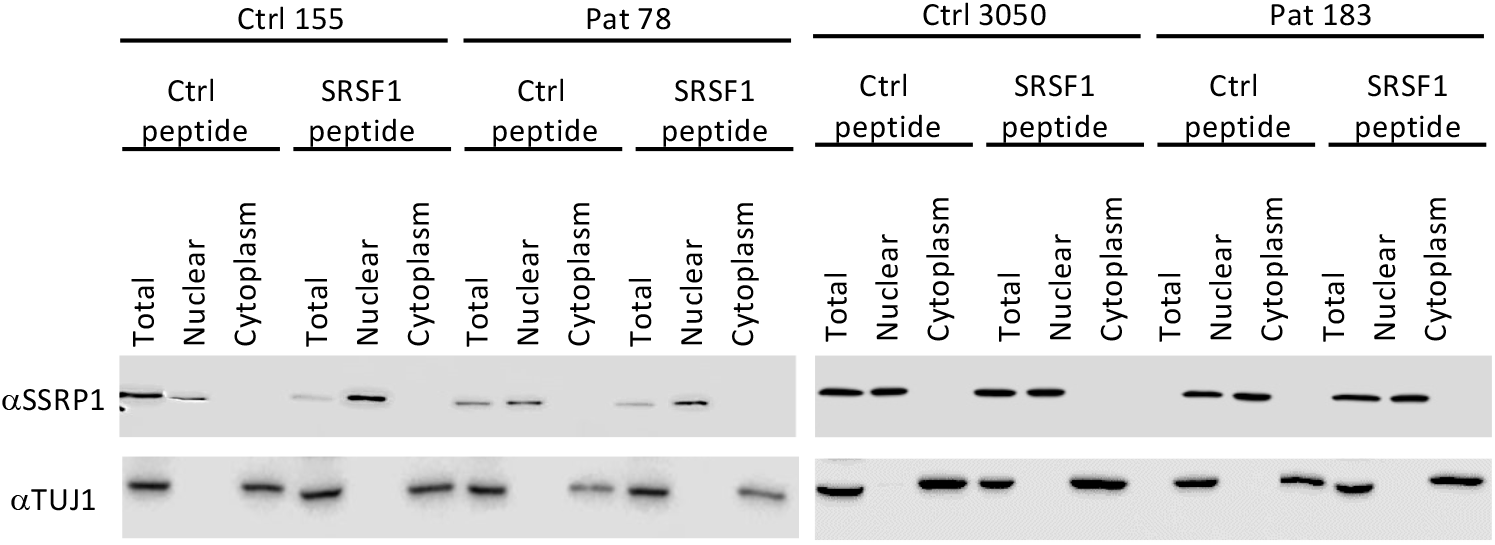
Preparation of total, nuclear and cytoplasmic fractions for quantifying the nuclear export of C9ORF72 wild type and repeat transcripts in healthy control and C9ORF72-ALS patient-derived neurons. Western blots of total, nuclear and cytoplasmic fractions isolated from patient-derived neurons treated for 72h with 5 mM of Ctrl or SRSF1 WT CPPs were: probed with nuclear chromatin-ass ociated SS RP1 factor and cytoplasmic pan neuronal marker beat III tubulin/TUJ 1.

**Table S1.**
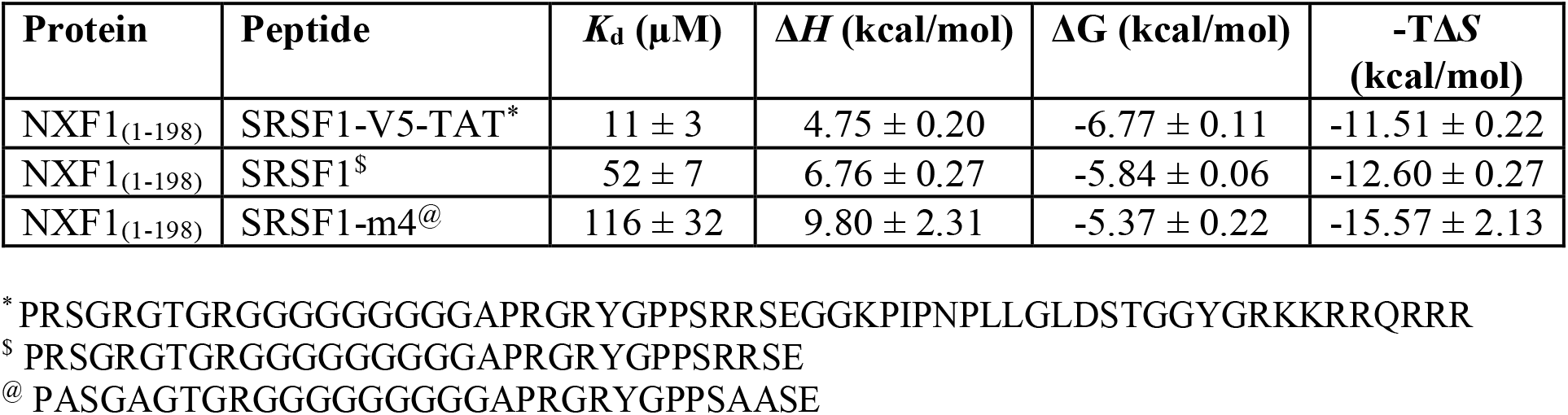
Thermodynamic parameters of NXF1 binding to SRSF1 WT and mutant peptides. Dissociation constants (*K*_d_) and thermodynamic parameters were derived from ITC data (Fig. S3) for NXF1 (aa1-198) in complex with SRSF1 WT-CPP (*), SRSF1 aa89-120 ($) and SRSF1 aa89-120 m4 (@).

**Table S2.** List and characteristics of patient-derived neurons used in this study.

| **Patient cell lines** | **Ethnicity** | **Gender** | **Cell type** | **Age at biopsy sample collection (years)** |
| --- | --- | --- | --- | --- |
| Ctrl 3050 | Caucasian | Male | Healthy control | 68 |
| Ctrl 155 | Caucasian | Male | Healthy control | 40 |
| Ctrl AG8620 | Caucasian | Female | Healthy control | 64 |
| C9-ALS p78 | Caucasian | Male | C9ORF72-ALS | 66 |
| C9-ALS p183 | Caucasian | Male | C9ORF72-ALS | 49 |
| C9-ALS p201 | Caucasian | Female | C9ORF72-ALS | 66 |

